# Computational modeling of DLBCL predicts response to BH3-mimetics

**DOI:** 10.1101/2023.02.01.526592

**Authors:** Ielyaas Cloete, Victoria M. Smith, Ross A. Jackson, Andrea Pepper, Chris Pepper, Meike Vogler, Martin J.S. Dyer, Simon Mitchell

## Abstract

In healthy cells, pro- and anti-apoptotic BCL2 family and BH3-only proteins are expressed in a delicate equilibrium. In contrast, this homeostasis is frequently perturbed in cancer cells due to the overexpression of anti-apoptotic BCL2 family proteins. Variability in the expression and sequestration of these proteins in Diffuse Large B cell Lymphoma (DLBCL) likely contributes to variability in response to BH3-mimetics. Successful deployment of BH3-mimetics in DLBCL requires reliable predictions of which lymphoma cells will respond. Here we show that a computational systems biology approach enables accurate prediction of the sensitivity of DLBCL cells to BH3-mimetics. We found that fractional killing of DLBCL, can be explained by cell-to-cell variability in the molecular abundances of signaling proteins. Importantly, by combining protein interaction data with a knowledge of genetic lesions in DLBCL cells, our *in silico* models accurately predict *in vitro* response to BH3-mimetics. Furthermore, through virtual DLBCL cells we predict synergistic combinations of BH3-mimetics, which we then experimentally validated. These results show that computational systems biology models of apoptotic signaling, when constrained by experimental data, can facilitate the rational assignment of efficacious targeted inhibitors in B cell malignancies, paving the way for development of more personalized approaches to treatment.

## Introduction

Apoptosis is a crucial component for the development of multicellular organisms and the functioning of the immune system. The BCL2 family of proteins are the principal regulators of mitochondrial-dependent apoptosis. This family of proteins consists of more than twenty-five members, and is further categorized into three groups based on their protein structure and function (Figure S1A and B): pro-apoptotic BCL2 proteins (BAX and BAK), anti-apoptotic BCL2 proteins (BCL2, BCL-xL, MCL1, etc.) and BCL2 homology domain 3 (BH3)-only proteins (BID, BIM, PUMA, NOXA, etc.).^1^ Prior to initiation of apoptosis, anti-apoptotic BCL2 proteins bind to BAX/BAK proteins at the mitochondrial outer membrane (MOM), impeding BAX/BAK oligomerization, which prevents mitochondrial outer membrane permeabilization (MOMP). Initiation of apoptosis leads to activation of BH3-only proteins, which either activate BAX/BAK directly through complex formation between BH3-only proteins and BAX/BAK or activate BAX/BAK indirectly by sequestering anti-apoptotic BCL2 proteins leading to the release of BAX/BAK from complexes containing the anti-apoptotic BCL2 proteins. BAX/BAK activation results in MOMP and subsequent apoptotic cell death.^2^

Avoidance of apoptosis is a hallmark of cancer, which in B cell lymphoma is often achieved through the upregulation of anti-apoptotic BCL2 proteins due to DNA translocation, gene amplification or constitutive activation of transcription factors that upregulate BCL2 family proteins such as nuclear factor kappa B (NF-κB).^3–5^ BCL2 dysregulation is commonly linked to chemoresistance and poor prognosis, and therefore represents a pathologically important biomarker and an attractive therapeutic target in B cell lymphoma.^6, 7^

ABT-199 (venetoclax), a BCL2 specific inhibitor, was first approved for treatment of chronic lymphocytic leukemia (CLL) and acute myeloid leukemia (AML),^8–10^ and has shown significant clinical activity in CLL regardless of genotype.^11^ However, in Diffuse Large B cell Lymphoma (DLBCL), responses to ABT-199 are less impressive, despite BCL2 overexpression in about 40% of cases.^8^ In a similar vein, BCL-xL is highly expressed in about 95% of DLBCL patient samples but only a proportion of DLBCL cell lines respond to BCL-xL inhibition.^12, 13^ Cell lines with comparably high levels of specific BCL2 family proteins frequently show different responses to BH3-mimetics that target those proteins.^13^ For example, there is no correlation between the abundance of MCL1 or BCL-xL and response to inhibitors that target these proteins^13^ and while BH3-profiling provides a functional measurement of the state of apoptotic signaling, actual responses to BH3-mimetics can differ from those predicted by BH3-profiling.^13, 14^ Collectively, these data indicate that the heterogeneous responses to BH3-mimetics in DLBCL are determined by the complex interactions between the BCL2 family of proteins and their binding partners.^12, 13^ Consequently, there is a need to develop better predictive tools to inform clinical decision making relating to optimal drug selection for individual patients.

Computational systems models can facilitate accurate prediction of how a molecular-scale signaling network will respond to perturbation with single cell and cell-population resolution.^15, 16^ Various aspects of the apoptotic signaling network have been encoded in computational models.^17–32^ However, DLBCL cells exhibit variable expression of multiple anti-apoptotic BCL2 proteins and show heterogeneous expressions of both pro-apoptotic BCL2 proteins and BH3-only proteins^12, 13^. Therefore, new computational models are required to capture the diverse abundances of BCL2 proteins implicated in DLBCL with their known interactional complexities to enable accurate prediction of responses to BH3-mimetics.

In this study, we aimed to establish virtual DLBCL cell lines generated from mechanistic computational models, informed by abundances of BCL2 family proteins. We aimed to use virtual cell lines to accurately predict, *in silico*, the experimentally-determined response of DLBCL cell lines to BH3-mimetics, and identify molecular and genetic determinants of treatment resistance. Finally, we sought to establish whether a computational systems biology approach can be used to rationally predict apoptotic responses in DLBCL by computationally identifying, and experimentally validating, novel synergistic combinations of BH3-mimetics. We aim to lay the foundation for a personalized medicine approach to targeting the spectrum of anti-apoptotic signaling found in B cell lymphoma.

## Materials and Methods

More detailed methods are provided in the supplemental Materials and Methods

### Experimental procedures

U2932 cells were maintained in RPMI-1640 media (21870076, Gibco; Life Technologies) with 10% fetal calf serum (10270-106, Gibco). Cells were plated at 4×10^5^ cells/mL and treated with ABT-199 and AZD5991 (Selleck Chemicals, Houston, TX)^33^ 24 hours before analysis using CellTiter-Glo viability assay (Promega, Madison, WI). Response was normalized to a DMSO control.

### BCL2 family signaling network model topology

The BCL2 family signaling network model was constructed from known protein interactions, requiring expression and degradation parameters for each protein,^29, 32, 34^ as well as binding rates for interacting proteins (Table S1).^29, 32^ The chosen set of interactions yields our BCL2 network topology (Figure 1).

**Figure 1.**
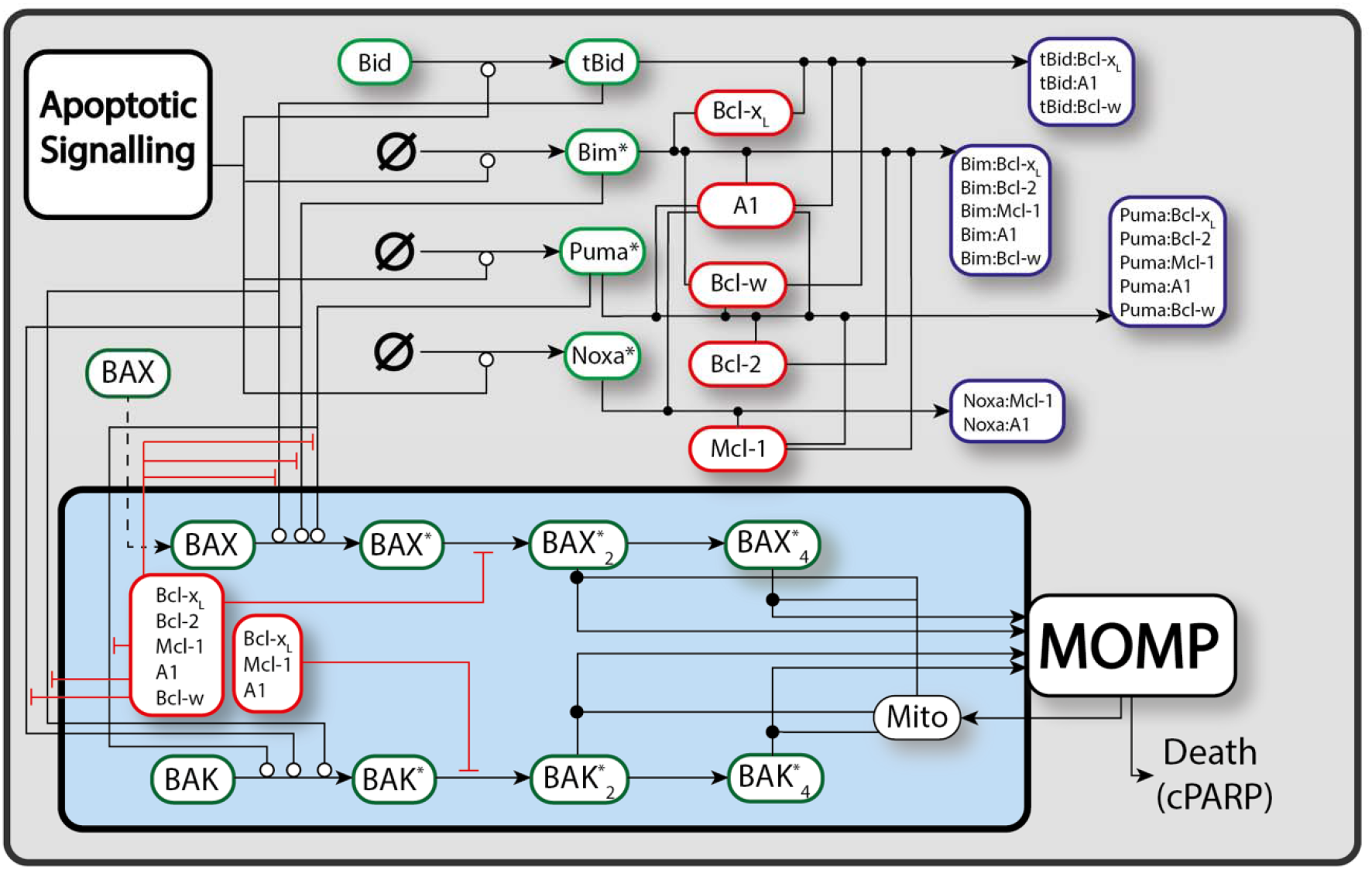
A schematic diagram of the apoptotic signaling network. Lines correspond to interactions between different species, with open circles, closed dots, and perpendicular lines denoting activation, binding, and inhibition/sequestration. Dashed lines correspond to the translocation of species. Some translocations are omitted as most BH3-only proteins and anti-apoptotic BCL2s are continuously trafficking between the cytoplasm and the MOM. Complete model details are provided in the supplemental modelling methodology and full code defining and running the model is available on GitHub (https://github.com/SiFTW/BH3Models).

### Model Construction

Model construction was achieved by building on previous models and incorporating experimentally measured BCL2 family protein expressions, binding affinities, kinetic rates, and knowledge of genetic lesions, see *Supplementary Text*.^13, 32, 34^ Parameter fitting was performed manually, informed by experimental data as described in the *Supplementary Text*.

Three *.csv* files were created for each virtual cell model, containing reactions between interacting proteins, the rate laws defining the reactions and a parameter file, respectively. Reactions between interacting proteins are governed by mass action kinetics, where interactions can be either simple binding and unbinding or binding/unbinding leading to activation of a new species. The three *.csv* files were used as inputs, to generate the system of Ordinary Differential Equations (ODEs) that defines our network model using custom python code (available: https://github.com/SiFTW/CSV2JuliaDiffEq).

### Solving Models/Running Simulations

The programatically generated model files were imported into Julia,^35^ and solved using the DifferentialEquations.jl package.^36^ Numerical simulations, initial conditions, solver options, and the code to generate all figures are available on GitHub (https://github.com/SiFTW/BH3Models).

### Simulating Cell-to-cell Variability

Cell populations of 100 cells were simulated with molecular cell-to-cell variability introduced in initial conditions. Initial conditions were distributed using a lognormal distribution with coefficient of variation (32%) defined by previous live-cell lineage tracking experiments in primary B cells^15^.

## Results

### A computational “unified-embedded-together” model enables exploration of differential sensitivities to BH3-mimetics

We hypothesized that the heterogeneous sensitivity of B cell lymphoma cells to BH3-mimetics is a predictable result of the state of the molecular signaling network in these cells. If true, an *in silico* computational model with sufficient detail and breadth would be able to predict responses, which could be validated experimentally. Combining the “embedded-together” and “unified” conceptual frameworks of apoptotic signaling (Figure S1A, SI supporting text),^37, 38^ and building upon established models of apoptotic signaling,^17–32^ we constructed a computational model capturing the complex interactions between BCL2 family proteins (Figure S1B) with appropriate granularity to simulate the effects of BH3-mimetics (Figure 1). Apoptosis-inducing signaling such as TRAIL or TNF receptor-proximal signaling is approximated as “apoptotic signaling” and kept fixed here, as the apoptosis-inducing signal we are studying is the addition of BH3-mimetics, which are explicitly modelled (Figure 1). We assume downstream effector-caspase induced biochemical and morphological alterations such as PARP cleavage is an inevitable consequence of MOMP. While the model does not explicitly include commonly mutated genes such as TP53 or MYC, these mutations can be simulated through their impact on kinetic rates in the signaling network.

### The experimentally-measured response to BH3-mimetics can be predicted by simulating a heterogeneous population of RC-K8 cells

To establish the feasibility of predicting the response of DLBCL cells to BH3-mimetics we first focused on the RC-K8 cell line, chosen due to lack of response to the BCL2 inhibitor ABT-199.^13^ We incorporated densitometry readings of BCL2 protein expression in RC-K8 cells measured by Western blotting into the newly established model in order to capture the expression profiles of BCL2 family proteins (Figure 2A, see SI supporting text).^13^ Comparing the predicted heterodimer abundances at steady state with co-immunoprecipitation experiments revealed a discrepancy (Figure S2).^13^ Given that previous work indicated that knowledge of the selective interactions between anti-apoptotic BCL2 proteins and BAX/BAK is a key determinant of sensitivity to BH3-mimetics, we incorporated the impact of complexes trafficking to subcellular localizations outside of the model’s scope by altering the degradation rates of complexes within the model’s signaling network (SI supporting text). The result was a virtual RC-K8 cell line with molecular network state that closely match expression profiles measured experimentally (Figure 2A and B). Simulating the impact of BCL-xL, BCL2 and MCL1 inhibition on MOMP in this virtual RC-K8 cell line resulted in strong induction of MOMP in response to BCL-xL inhibition, weak induction of MOMP in response to MCL1 inhibition and no response to BCL2 inhibition (Figure S3 A). This recapitulated the selective response of this line in experiments and indicated the model could predict the selective response of RC-K8 cells to BH3-mimetics.

**Figure 2.**
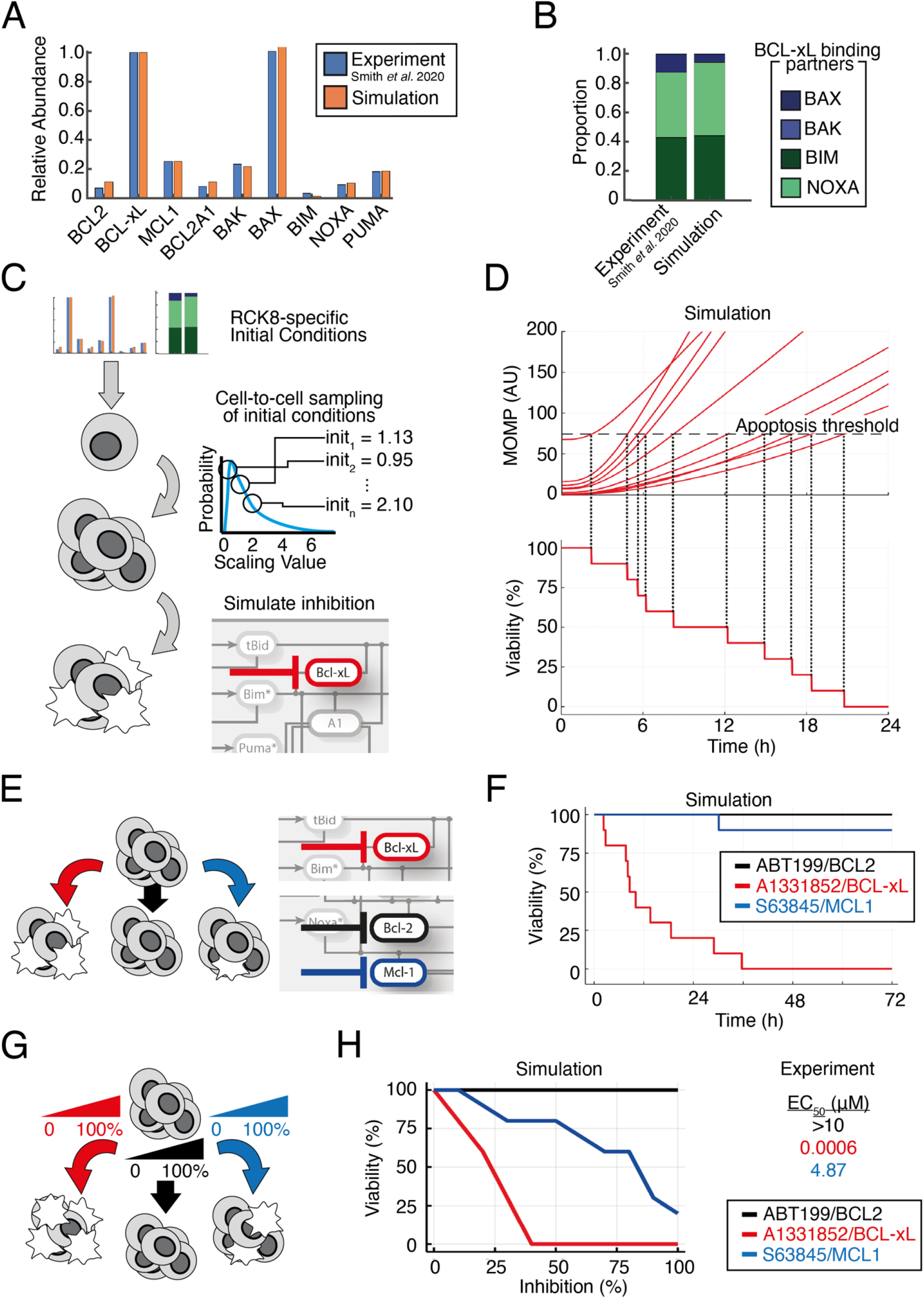
The experimentally-measured response to BH3-mimetics can be predicted by simulating a heterogeneous population of RC-K8 cells. **(A)** Comparison of the relative protein expression in the computational model to experimental data. Abundance of each protein is normalized to the most highly expressed anti-apoptotic BCL2 family protein and quantified from Smith *et al*.^13^ **(B)** Comparison of the proportion of pro-apoptotic and BH3-only BCL2 proteins bound to BCL-xL in the computational simulation compared to experimental data quantified from Smith *et al*.^13^ **(C)** Schematic of the method used to simulated BCL-xL inhibition in a heterogeneous cell population. From the RC-K8-speific parameterisation established in panels A and B initial conditions are independently sampled from a log-normal distribution to create heterogeneous cells with distinct starting states. Within all cells in the population the target protein (e.g. BCL-xL) is inhibited and the response to this perturbation recorded (see Methods). **(D)** Line graphs showing the simulated response to 50% BCL-xL inhibition in a heterogeneous RC-K8 cell population. A threshold of death (10% higher than is present within the naive population) is calculated. The time of death of in each cell is determined as the time this threshold is crossed (top panel). The percentage viability of the cell population can then be determined over time in response to BCL-xL inhibition (bottom panel). **(E)** Schematic showing that the process used to simulate BCL-xL inhibition in panels A to D is repeated for multiple BH3-mimetics. **(F)** Line graph of the simulated viability of the RC-K8 cell population in response to BCL2 inhibition (black, BCL-xL inhibition (red) and MCL1 inhibition (blue). The viability of the cell population is recorded to 72 hours to match experimental methods. **(G)** Schematic showing that the process used to simulate 50% inhibition in panels A to F is repeated for 10 distinct strengths of inhibition between 0 and 100% to enable comparison to dose-response experiments. **(H)** Line graphs showing the simulated percentage of the RC-K8 cell population viable at each percentage inhibition of the indicated target protein. This prediction can be compared to experimentally measured EC_50_ values, right.^13^

As a fixed and reproducible fraction of the DLBCL cells undergo apoptosis in response to a given dose of BH3-mimetic,^13^ we hypothesized that this may result from molecular cell-to-cell heterogeneity within the cell population. The cell-to-cell variability in the abundance of signaling components in B cells has been previously quantified through combined lineage tracing and computational modelling, and was found to predictably explain fixed proportions of primary B cells undergoing distinct fates such as apoptosis, mitosis and differentiation in response to antigenic stimulation.^15,16^ We therefore converted these simulations of a single average cell-population response, into simulations representing heterogeneous populations of single cells by sampling initial conditions as described previously (Figure 2C).^15^ Simulating 50% inhibition of BCL-xL revealed that distributed initial conditions were sufficient to create heterogeneity in the timing that MOMP increased within a simulated cell population (Figure 2D). We considered an individual cell to have died in response to BCL-xL when MOMP exceeded a threshold of 10% above the level of MOMP in the pre-treatment population and, in this way, we were able to simulate viability over time in response to BH3-mimetics in the virtual RC-K8 cells (Figure 2D). Extending this approach to the effect of other BH3-mimetics revealed specific timings of apoptosis within the cell population, and the proportion of cells undergoing apoptosis in response to 50% inhibition of BCL2, BCL-xL and MCL1, with the largest proportion of cell death occurring rapidly in simulations of RC-K8 cells following BCL-xL inhibition (Figure 2E and F). Simulating 0-100% inhibition of BCL2, BCL-xL and MCL1 within a heterogeneous cell population enabled predictions of the proportion of the RC-K8 cells that would be viable at 72 hours (Figure 2G and H); these predictions were comparable to experimentally determined EC_50_ values; the concentration of a drug required to produce 50% of its maximal effect (Figure 2H).^13^ We find that this systems biology approach predicts that RC-K8s have low EC_50_ BCL-xL inhibitors, high EC_50_ for MCL1 inhibitors, and that the EC_50_ for BCL2 inhibitors will not be reached even at 100% inhibition of BCL2 (Figure 2H). This is in strong agreement with experimentally determined EC_50_ values (Figure 2H right), and independent of the specific threshold of MOMP activity chosen to trigger apoptosis (Figure S3B).

### Virtual lymphoma simulations can predict response to BH3-mimetics

As we found a systems biology approach could predict the response of RC-K8 cells to BH3-mimetics, we expanded the approach to a library of six DLBCL lines (RIVA, U2932, RC-K8, SUDHL8, SUDHL10, and U2946), chosen to capture the diversity of responses to BH3-mimetics seen within both patient samples and cell lines.^13^ We incorporated the heterogeneous expression of BCL2 family proteins and their heterodimerization profiles, measured by immunoprecipitation and co-immunoprecipitation, in the same way as we did for RC-K8 cells (Figure 2).^13^ The result was a library of virtual DLBCL cell lines that accurately captured the abundance and binding partners of BCL2 family proteins (Figure 3A). Simulating the response of the library of DLBCL lines to BH3-mimetics targeting BCL2, BCL-xL and MCL1 predicted highly heterogeneous responses (Figure 3B). Simulations predicted that RIVA cells would only respond to BCL2 inhibition, while U2932 cells were predicted to be broadly resistant to all BH3-mimetics (Figure 3B). Comparing this prediction to experimentally determined EC_50_ values showed that the model had correctly predicted that RIVA cells only responded to BCL2-targeting ABT-199, while the predicted lack of response in U2932 cells was confirmed by the high (micromolar) EC_50_ values of U2932 cells when challenged with all three BH3-mimetics (Figure 3B). Both RC-K8 cells and SUDHL8 cells were predicted to be most sensitive to inhibition of BCL-xL, with both lines also responding to MCL1 inhibition (at intermediate doses in SUDHL8 cells, and high doses in RC-K8 cells) (Figure 3B). Comparing these predictions to experimentally determined EC_50_ values revealed that these computational predictions closely match experimental measurements (Figure 3B). Simulation of SUDHL10 and U2946 cell lines predicted that these lines would only respond to inhibition of MCL1, which was validated by experimental measurements (Figure 3B). Taken together, our virtual lymphoma simulations accurately predicted experimentally measured responses to BH3-mimetics.

**Figure 3.**
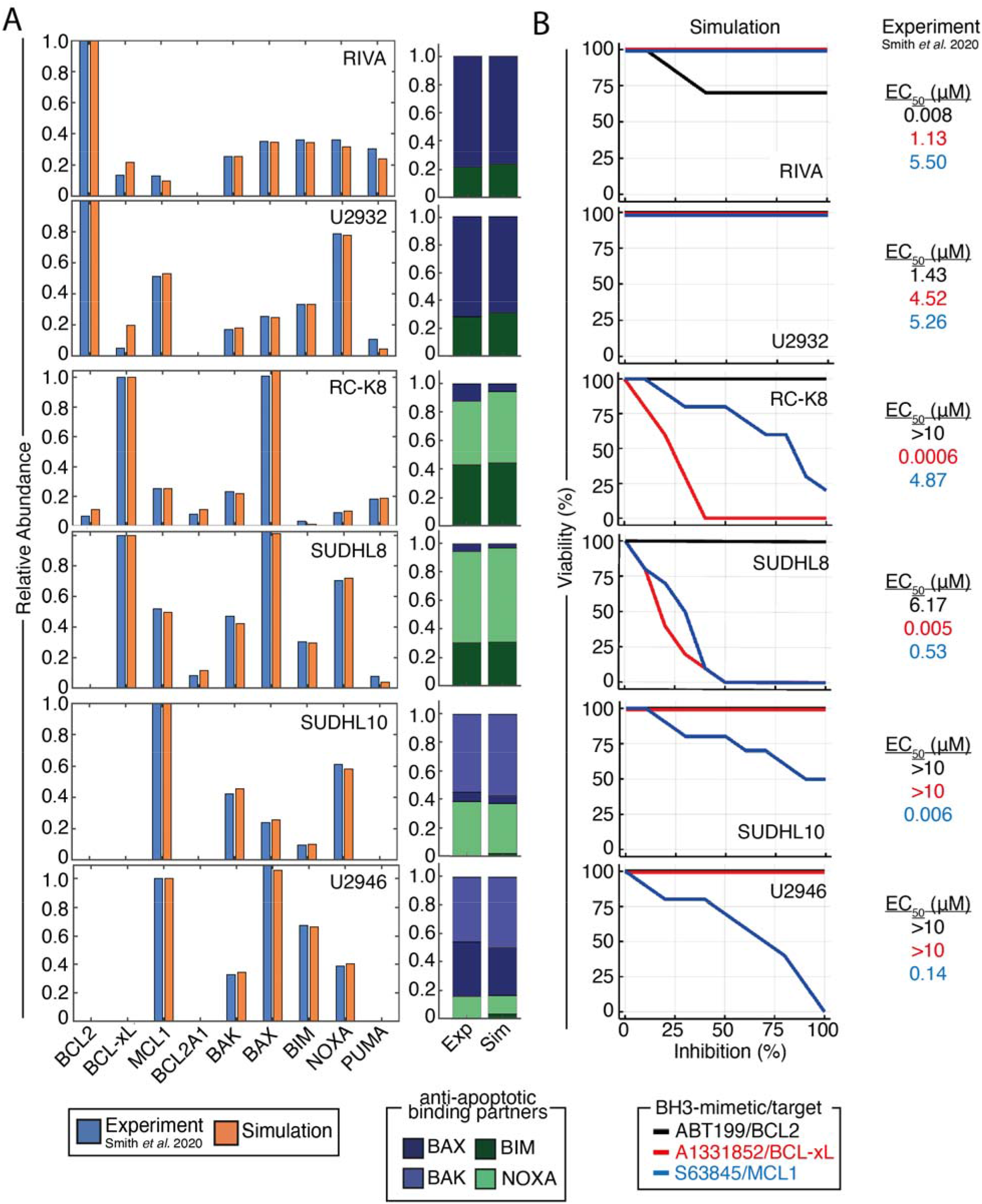
Cell line-specific computational models recapitulate experimentally measured protein expression and heterodimerisation profiles, and enable accurate prediction of the optimal BH3-mimetic for each cell line. **(A)** Right: Comparison of the relative protein expression in the computational model to experimental data. Abundance of each protein is normalized to the most highly expressed anti-apoptotic BCL2 family protein and quantified from Smith *et al*.^13^ Left: Comparison of the proportion of pro-apoptotic and BH3-only BCL2 proteins bound to the dependent anti-apoptotic BCL2 protein in each line in the computational simulation compared to experimental data. **(B)** Model simulations (left) of cell population viability (%) in response to 10 different strengths of BH3-mimetics compared to experimentally-measured EC_50_ values.

### Considering genetic lesions in virtual lymphoma can improve accuracy of simulations

While the computational model could accurately assign the right drug to the right DLBCL cell lines informed by protein-expression data alone, some quantitative discrepancies between computationally predicted responses and experimentally measured EC_50_ values suggests that additional mechanisms may contribute to selective responses. Genetic lesions, such as MYC translocations and p53 mutations, present in all modeled cell lines except RC-K8 cells,^39^ can have numerous potential consequences, and the most functionally significant impact of these mutations is likely on the apoptotic network state. We sought to predict which of these potential impacts was functionally significant in controlling the response to BH3-mimetics by computationally identifying which effect improved the match between the computational predictions and experimental validation.

RIVA cells were more sensitive to BCL2 inhibition than predicted from protein data alone (Figure 3B). Simulating the impact of BCL2 gene amplification and MYC translocation (resulting in elevated BAX expression),^40^ increased the sensitivity of virtual RIVA cells to ABT-199 indicating an important role for these mechanisms in modulating the response to BH3-mimetics (Figure S4A and B). Incorporating the presence of a MYC-overexpressing subclone in U2932 cells, which resulted in increased BIM and BAX expression, explained the response of this line to high doses of ABT-199 (Figure S4A and B). In RC-K8 cells the magnitude of the difference in response to A1331852 and S63845 was underestimated by our simulations (Figure 3B). The match between simulation and experiment in both lines was improved by decreasing the abundance of MCL1. Biologically, this could be mediated by the truncated p300 expressed in RC-K8 cells, which reduces the acetylation of MCL1 thereby decreasing MCL1 protein stability.^41^ In SUDHL8 cells, mutated TP53 may cause the reduced gene expression of MCL1 (Figure S4A and B).^42^ While the simulation accurately predicted the response of SUDHL10 and U2946 cells to the MCL1 inhibitor S63845, the simulation only predicted apoptosis at high doses of the inhibitor (Figure 3). MYC translocation in SUDHL10 may increase expression of MCL1, BIM and NOXA resulting in increased sensitivity to inhibition of MCL1, while mutated TP53 in SUDHL10 cells may decrease the affinity of MCL1 for p53 protein, increasing the binding between MCL1 and BIM (Figure S4A and B).^43^ Taken together these data show that by comparing computational predictions with experiment results, and iteratively improving the match between the two, we can narrow down the plethora of potential effects of genetic lesions to those that are predicted to be functionally significant. By iteratively improving the model in this way, the correlation between the predicted response from simulations and experiments across the library of virtual cell lines substantially improved (R^2^ from 0.38 to 0.67, Figure S4C and D). This suggests that while the optimum BH3-mimetic can be reliably identified from protein data alone (Figure 3), once genetic lesions are considered, the virtual cell lines can quantitatively predict experimentally determined EC_50_s in virtual cell lines (Figure S4D).

### Inherent resistance to BH3-mimetics in cellular sub-populations is the predictable result of cell-to-cell molecular variability

As fractional killing of a cell line can be explained as the predictable result of inherent molecular heterogeneity between cells (Figure 2), we sought to use simulations to predict the molecular determinants of inherent resistance to BH3-mimetics. Simulating a dose of BH3-mimetics that causes 50% reduction in viability and then analyzing the starting state of cells revealed statistically significant differences (p<0.05) in the predicted molecular network state of cells that would undergo apoptosis in response to the inhibitor when compared to those that would be resistant to the treatment (Figure S5-12). As expected, within a population of RIVA (Figure S5), U2932 (Figure S6), SUDHL10 (Figure S7) and U2946 cells (Figure S8) responding to BCL2 inhibition, the treatment-sensitive cells had higher abundances of pro-apoptotic and lower anti-apoptotic BCL2 proteins prior to the drug being applied.

Furthermore, cells that undergo apoptosis were predicted to have fewer complexes between anti-apoptotic BCL2 proteins and BH3-only proteins (Figure S5-6). In RC-K8 cells responding to BCL-xL inhibition (Figure S9) and MCL1 inhibition (Figure S10), the cells that undergo apoptosis expressed more pro-apoptotic BCL2 proteins, more BH3-only proteins, and more complexes between pro-apoptotic BCL2 and BH3-only proteins. Intriguingly, cells susceptible to BCL-xL and MCL1 inhibition had significantly more pro-apoptotic proteins bound to the mitochondria (Figure S9 and 10).

While BCL2A1 was not predicted to contribute to the inherent treatment-resistant cell population in some cell lines (U2932, RIVA, SUDHL10, and U2946), in others (RC-K8 and SUDHL8) the BCL-xL resistant cell populations were predicted to be significantly higher in BCL2A1 (Figure S9-12). Interestingly, in SUDHL8 cells responding to MCL1 inhibition a significant role was predicted for BCL2A1, while this was not seen in RC-K8 cells (Figure S10 and S12).

Virtual cell lines reveal that inherent cell-to-cell variability in the sequestration of anti-apoptotic BCL2 proteins, and subcellular localization of pro-apoptotic complexes, can create an inherent treatment resistant cell population. Importantly, we found that the mechanisms of inherent treatment resistance were predicted to be diverse between different cell lines and BH3-mimetics.

### Synergistic combinations of BH3-mimetics can be computationally identified and experimentally validated

To test the ability of this systems biology approach to enable rational targeting of DLBCL we simulated combinations of BH3-mimetics and identified a number of synergistic combinations (Figure S13). In BCL-xL dependent cell lines, RC-K8 (Figure S13C) and SUDHL8 (Figure S13D), synergy was predicted between BCL-xL and MCL1 inhibitors.

Additionally, in RC-K8 cells, the model predicted synergy between BCL-xL and BCL2 inhibition. The exquisite sensitivity of RC-K8 cells to A1331852 monotherapy (BCL-xL -targeting BH3-mimetc) meant that experimentally testing this predicted synergy would be challenging and unlikely to identify therapeutically significant combination therapies.

Simulations in virtual U2932 cells predicted no response to MCL1 inhibition alone. However, adding MCL1 inhibition in the context of BCL2 inhibition was predicted to synergistically induce apoptosis (Figure 4A, left). Predicting the impact of combining BCL2 and MCL1 inhibition across a broad range of doses indicated that it should be possible to experimentally test this prediction (Figure 4B left, average Bliss synergy score: 15.31). The model prediction of synergy between BCL2 and MCL1 inhibitors in U2932 was experimentally tested by treating U2932 cells with ABT-199, AZD5991 and the combination in equimolar concentrations to recapitulate the computational predictions. In striking concordance with the computational prediction, U2932 cells were insensitive to the MCL1-specific monotherapy (AZD5991) but in combination with the BCL2-specific BH3-mimetic, ABT-199, showed synergistic induction of apoptosis (Figure 4A, right). Extending this analysis to all combinations of doses in both the computational model and experimental system (Figure 4B, Figure S14) confirmed the combination of BH3-mimetics was synergistic (average Bliss synergy score: 23.96). While 50% inhibition of MCL1 and BCL2 was predicted to kill 0% and 50% of cells, respectively, the combination was predicted to result in a 70% reduction in viability. This was confirmed by our experimental findings, which showed that 100 nM of AZD5991 and ABT-199 killed approximately 0% and 40% of cells, respectively while combining these two agents at the same concentrations induced over 70% cell death (Figure 4B). Computationally identifying and experimentally validating synergistic combinations of BH3-mimetics demonstrates that a systems biology approach has the potential to enable rational assignment of efficacious targeted therapies in DLBCL.

**Figure 4.**
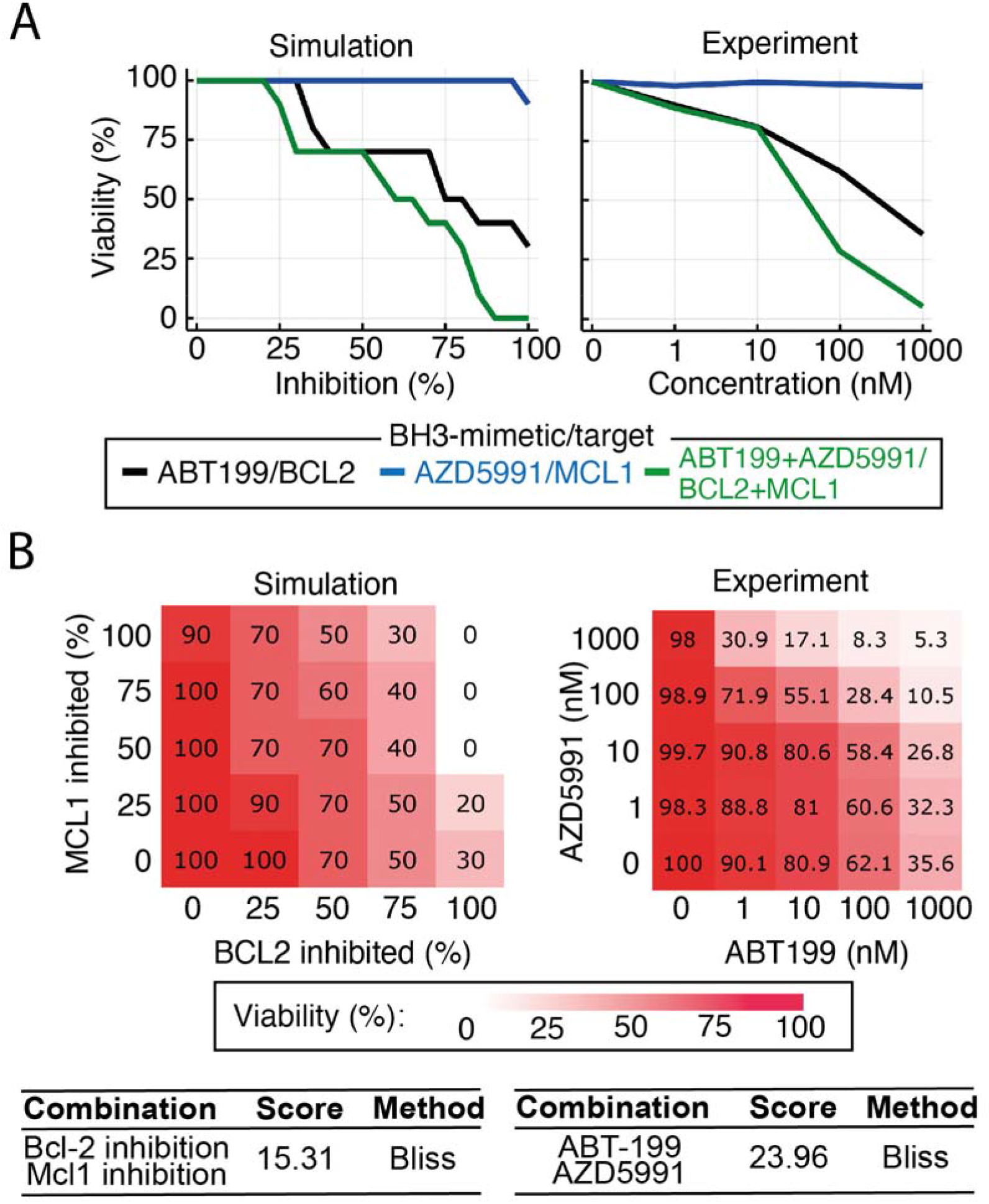
Synergistic interactions between BH3-mimetics can be computationally identified and then experimentally validated. **(A)** (Left panel) Model simulation predicting synergy between BCL2 and MCL1 inhibition in U-2932 cells (green curve). The model predicts a moderate response to BCL2 inhibition alone (black curve), no response to MCL1 inhibition alone (blue curve) but a greater than additive response when MCL1 and BCL2 inhibition is combined at equal levels. (Right panel) Experimental data validates the model prediction, demonstrating synergy between the BCL2-specific inhibitor, ABT-199, and the MCL1-specific inhibitor, AZD5991. The combination was tested at equimolar concentrations. **(B)** Complete doseresponse matrices showing the impact on cell viability of all combinations of simulated inhibition strengths (left) and experimentally measured BH3-mimetic doses (right) for BCL2 inhibition (ABT-199) and MCL1 inhibition (AZD5991). Viability was calculated at 72 hours for both computational models and experimental measurements. Bliss synergy scores of the model simulation (left panel) and experimental data (right panel), calculated using SynergyFinder software.^49^

## Discussion

In this study we created a novel computational model of apoptosis and, using experimental data, we established a library of virtual DLBCL cell lines that could reliably predict experimentally measured responses to BH3-mimetics. Importantly, this systems biology approach was also capable of identifying novel synergistic combinations of inhibitors, which we then experimentally validated. The success of our approach when using a library of heterogeneous cell lines indicates that it might be feasible to expand this approach to patient-derived samples, which may form the foundation of a personalized treatment approach in DLBCL.

While the model constructed here was based on B cell lymphoma, many of these apoptotic interactions are conserved in other cell types both in health and malignancies.^26, 44^ To facilitate translating this approach to the challenges associated with the rational assignment of targeted inhibitors in other hematological and solid malignancies, we have made the complete library of virtual cell lines freely available (https://github.com/SiFTW/BH3Models).

One strength of computational modeling is the ability to incorporate and test the consistency of multiple modalities of data and assimilate them into a model that encapsulates the current state of knowledge for a given system. While we acknowledge that immunoprecipitation and coimmunoprecipitation blots provide semi-quantitative data, the ability of this data to create models capable of accurate predictions across a library of cell lines and BH3-mimetics indicates that our modelling approach is robust to the inherent variability of such data. While proteomic data may provide more precise quantification, this data critically lacks information on binding partners that were found to be key to determining response. Given the challenges of obtaining immunoprecipitation data for each patient, future work to identify surrogate biomarkers that provide clinically-obtainable measurements for the underlying network state will be required. Modelling results can be used to inform statistical regression to identify the most informative parameters that can be experimentally measured and perturbed. Such approaches have been successfully experimentally validated in B cells.^15^

The model utilized here is a sub-network of the apoptotic signaling network. As such, the model only accounts for the mutational and molecular heterogeneity in a proportion of the important signaling networks implicated in B cell malignancies. To assess the impact of dysregulation of multiple signaling networks (including upstream receptor proximal signaling, NF-κB regulation, cell cycle and differentiation), and treatment modalities targeting these networks, future work will necessarily require more comprehensive models. Established multi-scale B cell models containing these signaling networks and linking them to cell fates have demonstrated utility in predicting the emergent impact of genetic events and inhibitors on cell population phenotypes.^15, 16^ Re-purposing these models for lymphoma, and incorporating the insight generated here, may enable rational targeting of additional therapies and combinations of therapies that target these signaling networks (such as ibrutinib^45^, idelalisib^46^, and copanlisib^47, 48^).

While we have tested many of the computationally generated predictions in this study in order to validate our models, many intriguing predictions remain as avenues for future work. For example, predictions of which cells will survive BH3-mimetic treatment may provide insight into how combination therapies or pre-treatments might be deployed to optimize the molecular network state for sensitivity to BH3-mimetics. It is likely that treatment-resistant cells are vulnerable to alternative targets as each apoptotic network state we investigated had both resistances and vulnerabilities. The cell lines we investigated covered a broad range of expression of BCL2 family member expression, a broad variety of protein-protein interaction states, multiple cell-of-origin classifications, and common mutational events (Figure 3 and Figure S4). However, the existence of B cell lymphoma lines (such as DLBCL line HBL-1, and primary mediastinal B cell lymphoma line such as Karpas-1106) that do not respond well to any BH3-mimetic indicates that additional cellular archetypes exist.^13^ Identifying therapeutic vulnerabilities in the network state of these broadly resistant cells may help target the most challenging cases. However, as these lines are in the minority, the primary challenge in adopting more targeted therapies in B cell lymphoma appears to be getting the right combination of drugs that are already available into the right patients. Systems biology modeling may enable this and by modeling individual patients unlock the potential of personalized medicine.

## Data sharing statement

All code used in the current study are available in the GitHub repository, https://github.com/SiFTW/CSV2JuliaDiffEq.

## Acknowledgements

The authors would like to thank Eleanor Jayawant and Timothy Chevassut for critical reading of the manuscript, and the following funders. SM: UKRI Future Leaders Fellowship (MR/T041889/1), leukaemia UK John Goldman Fellowship (2020/JGF/003) and a Beat:Cancer research grant. AP: MRC Research Grant (MR/V009095/1). MV: Else Kröner-Fresenius-Stiftung. VMS, RAJ and MJSD: This work was supported by funds from the Scott-Waudby Trust, and the Kay Kendall Leukemia Fund.

