## Supplementary File for "Computational modeling of DLBCL predicts response to BH3-mimetics"

### Supporting Information Text

#### The conceptual frameworks used in modelling

Multiple conceptual models/frameworks of the complex interaction between BCL2 family members have been proposed, including the direct activation model, the displacement or de-repression model, embedded-together model and the unified model.[1-11] A review of these conceptual frameworks is given in Shamas-Din *et al.*[12] We consider both the embedded-together and unified models, which introduce a role for the mitochondrial membrane, as MOMP does not ensue until BAX and BAK achieve their active conformation in the membrane. Furthermore, in both these models, anti-apoptotic BCL2 proteins sequester BH3-only proteins and pro-apoptotic BCL2 proteins. However, a distinguishing feature of the unified model is that inhibition of apoptosis via sequestration of pro-apoptotic BAX and BAK is more efficacious than inhibition via sequestration of BH3-only proteins. To capture the range of potential interactions in B cell lymphoma cells, we encoded features from both conceptual models, along with previously published models, and recently published experimental data into our computational model (Figure 1A).[2, 6, 13-28] Briefly, we consider inactivator or sensitizer BH3-only proteins to interact exclusively with anti-apoptotic BCL2 proteins impeding the translocation of anti-apoptotic BCL2 to the mitochondrial membrane and the subsequent interaction of anti-apoptotic BCL2's with pro-apoptotic BAX and BAK and activator BH3-only proteins.[4, 7, 29] While we consider activator BH3-only proteins to perform the dual function of directly activating pro-apoptotic BCL2 proteins, initiating membrane permeabilization and also sequestering anti-apoptotic BCL2 proteins, inhibiting their interaction with pro-apoptotic BCL2 proteins.[10, 11]

Having established the conceptual framework, we then encoded knowledge of the selective interactions between BCL2 family proteins. We considered that all anti-apoptotic BCL2 proteins interact with pro-apoptotic BAX through their conserved BH3 binding groove, while, among the anti-apoptotic BCL2 family, only BCL-xL, MCL1 and BCL2A1 interact with pro-apoptotic BAK, also through their conserved BH3 binding groove.[5, 8, 11, 30-33] We considered that BH3-only proteins bind to anti-apoptotic BCL2 proteins in the same binding groove but with greater selectivity for binding partners. Bim and Puma interact with all anti-apoptotic BCL2 proteins, while Bid or its truncated variant tBid binds to BCL-xL, BCL2A1 and BCL-w, and Noxa binds selectively to MCL1 and BCL2A1. Furthermore, BID (tBID), BIM and PUMA but not NOXA are known to activate both BAX and BAK, resulting in BAX and BAK binding to the mitochondrial membrane and MOMP (Figure 1 and S1).[1, 11, 12, 30-34]

#### Supplementary Modelling Methodology

The BCL2 family of proteins is classified into three groups: pro-apoptotic BCL2 proteins, which interact directly with the mitochondrial outer membrane, permeabilizing the membrane; BH3-only proteins (BH3s), which directly or indirectly activate pro-apoptotic BCL2 proteins; and anti-apoptotic BCL2 proteins, which directly or indirectly inhibit pro-apoptotic activation and pore formation.[12] Adding an extra layer of complexity, interactions of BCL2 proteins are regulated by which family members are present, their relative binding affinities and concentrations, as well as their spatial localization.

In this work, inactivator or sensitizer BH3s interact exclusively with anti-apoptotic BCL2 proteins impeding the translocation of anti-apoptotic BCL2 to the mitochondrial membrane and the subsequent interaction of anti-apoptotic BCL2s with pro-apoptotic BAX and BAK and activator BH3s.[4, 7, 29] Activator BH3s perform the dual function of directly activating pro-apoptotic BCL2 proteins, initiating membrane permeabilization and also sequestering anti-apoptotic BCL2 proteins, inhibiting their interaction with pro-apoptotic BCL2 proteins.[10, 11]

In Figure 1, Apoptotic signaling acts as a proxy for the extrinsic apoptotic stimuli and the caspase network and, thus, activates the BH3s in the model. The model consists of two compartments, the cytoplasm (grey shaded region) and the mitochondrial outer membrane (MOM) (blue shaded region). Lines in the diagram denote interactions between species, with open circles corresponding to an activating interaction, closed dots corresponding to a binding interaction and perpendicular lines corresponding to inhibition/sequestration.

Experimental studies have demonstrated how, when exposed to extrinsic stimuli, cells undergo a protracted and variable delay preceding caspase activation (preceding BH3-only activation in this model), but a swift evolution to MOMP once activation is initiated.[35, 36] Similar “variable-delay, snap-action” switching behavior seen experimentally has been observed in previous models of apoptosis.[22, 24, 28] Our variable of interest in the model is MOMP, which is a proxy for the proportion of PARP that has been cleaved and, therefore, also a proxy for cell death. Figure S3 (A) shows the result of our model simulation for the level of MOMP as a function of time for different apoptotic signaling doses (labelled Signal in the figure). As seen in both experimental data and computational models, the rate of increase of MOMP increases with increasing doses of the apoptotic signal. However, MOMP saturates at a higher level in our model as the apoptotic signal increases. In previous models, the proportion of cleaved PARP (cPARP) is used as the response variable and appears to reach the same saturation level for different doses of extrinsic death. One possible explanation is that the positive feedback from MOMP via caspase-3 and caspase-8 results in a threshold level of MOMP generating more MOMP. Since we have omitted the caspase network in our model, it is reasonable to assume that our system is equivalent to previous systems, including the positive feedback mechanism. Thus, MOMP is a legitimate proxy for cPARP and cell death.

BH3-mimetics function to sequester anti-apoptotic BCL2 proteins, releasing pro-apoptotic BAX and BAK. However, in our model simulations, we model BH3-mimetic activity as anti-apoptotic BCL2 protein inhibition. Figure S3 (A) shows the response to anti-apoptotic BCL2 family inhibitors our model (20 and 50% inhibition).

**Parameter Fitting**
Expression and degradation parameters and binding and unbinding rates for the network model were taken from published results including computational models developed for other biological contexts.[24, 28, 37] Cell line-specific BCL2 protein expression parameters were manually adjusted to recapitulate experimentally measured protein expression profiles in each cell line.[38] Experimentally measured expression and dimerization profiles were quantified using ImageJ.[39] The emergent heterodimerization profiles were compared to experimentally measured dimerization profiles.[38] To address discrepancies between simulated heterodimer profiles and experimentally measured heterodimerization profiles, we assume that BCL2 protein homodimers and heterodimers are localized in multiple locations across the cell, including the endoplasmic reticulum, the Golgi apparatus and the nucleus, and that biochemical measurements of protein expression will account for all the proteins regardless of location. However, we only model BCL2 proteins in the cytoplasm and MOM-bound. Therefore, we accounted for complexes trafficked to subcellular localizations outside of the model's scope by manually altering the degradation rates of protein complexes within the model's signaling network. Upon establishing accurate protein expression and binding profiles for each line’s parameters were kept consistent for all subsequent simulations, other than those influenced by genetic lesions (see Table S1).

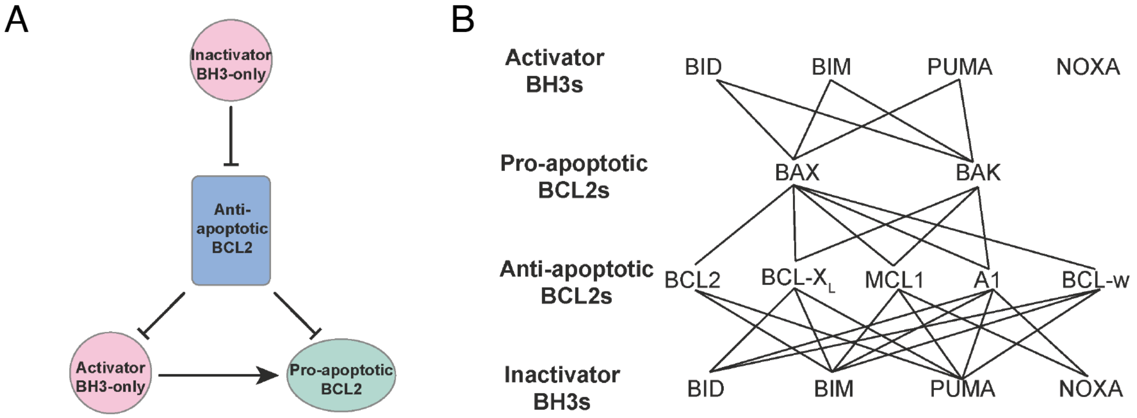


**Figure S1.** A unified-embedded-together model of apoptosis. (A) Schematic diagram of the unified-embedded-together model. (B) Graph of the BCL2 family of protein interactions. Nodes correspond to BCL2 family members, while links indicate interactions between corresponding proteins.

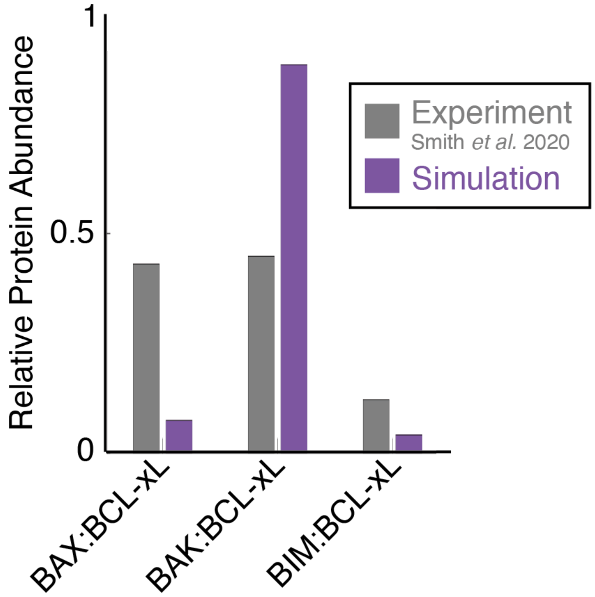


**Figure S2.** Heterodimerization profile in RC-K8 cells that emerges from protein expression profile in Figure 1A in both experimental data (grey[38]) and model simulations (purple). Expression values are normalized to show proportion of BCL-xL bound to BAX, BAK and BIM.[38]

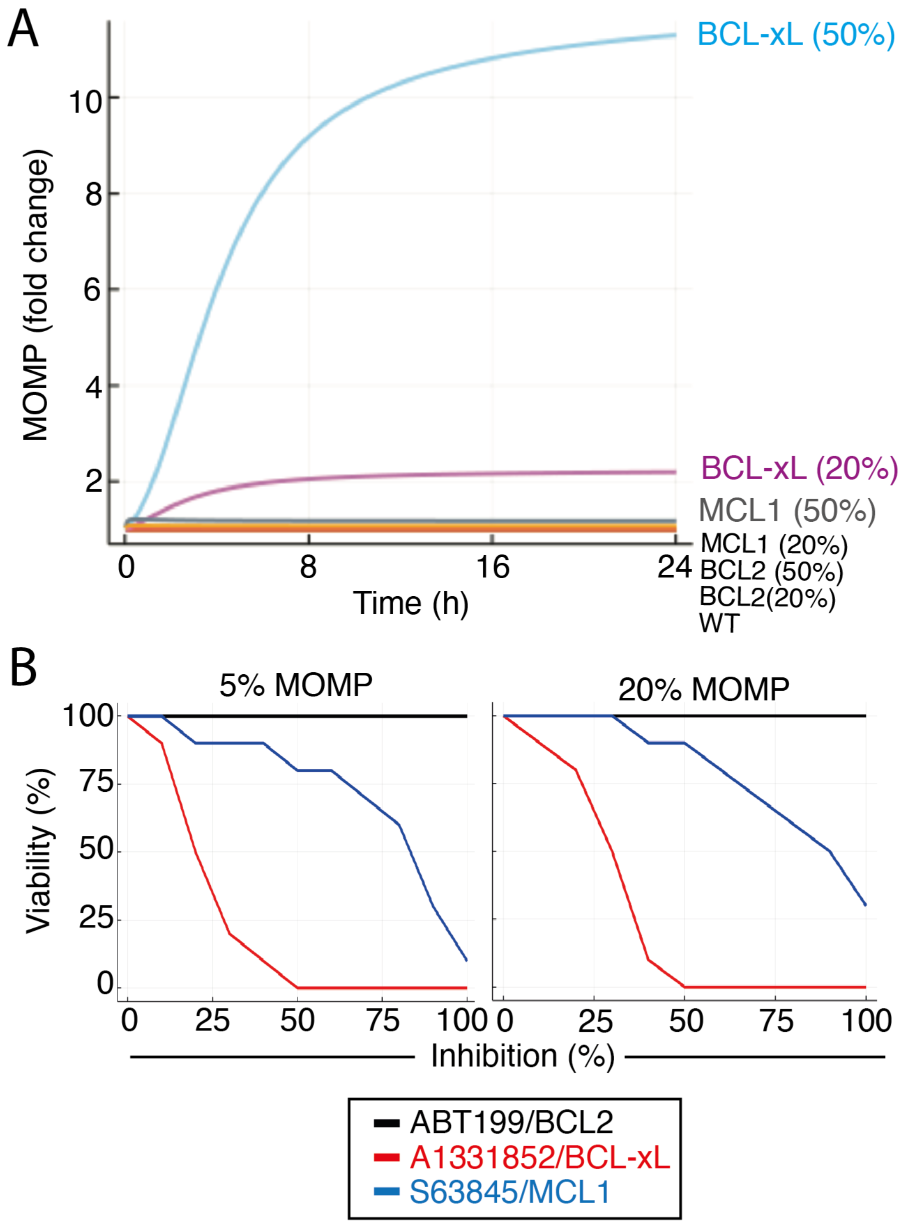


**Figure S3.** (A) RC-K8 specific response to BH3-mimetics. The model simulation displays the level of MOMP achieved in response to the indicated level of inhibition of the indicated BCL2 family protein. The level of MOMP is measured as a fold change from the basal level of MOMP (MOMP level before administering the BCL2 inhibitor). (B) The viability (%) in response to inhibition of the indicated BCL2 family proteins for the respective virtual cell lines with MOMP threshold decreased 2x from the value used in Figure 2 (left, 5% MOMP inhibition), and increase 2x from the value used in Figure 2 (right, 5% MOMP inhibition).

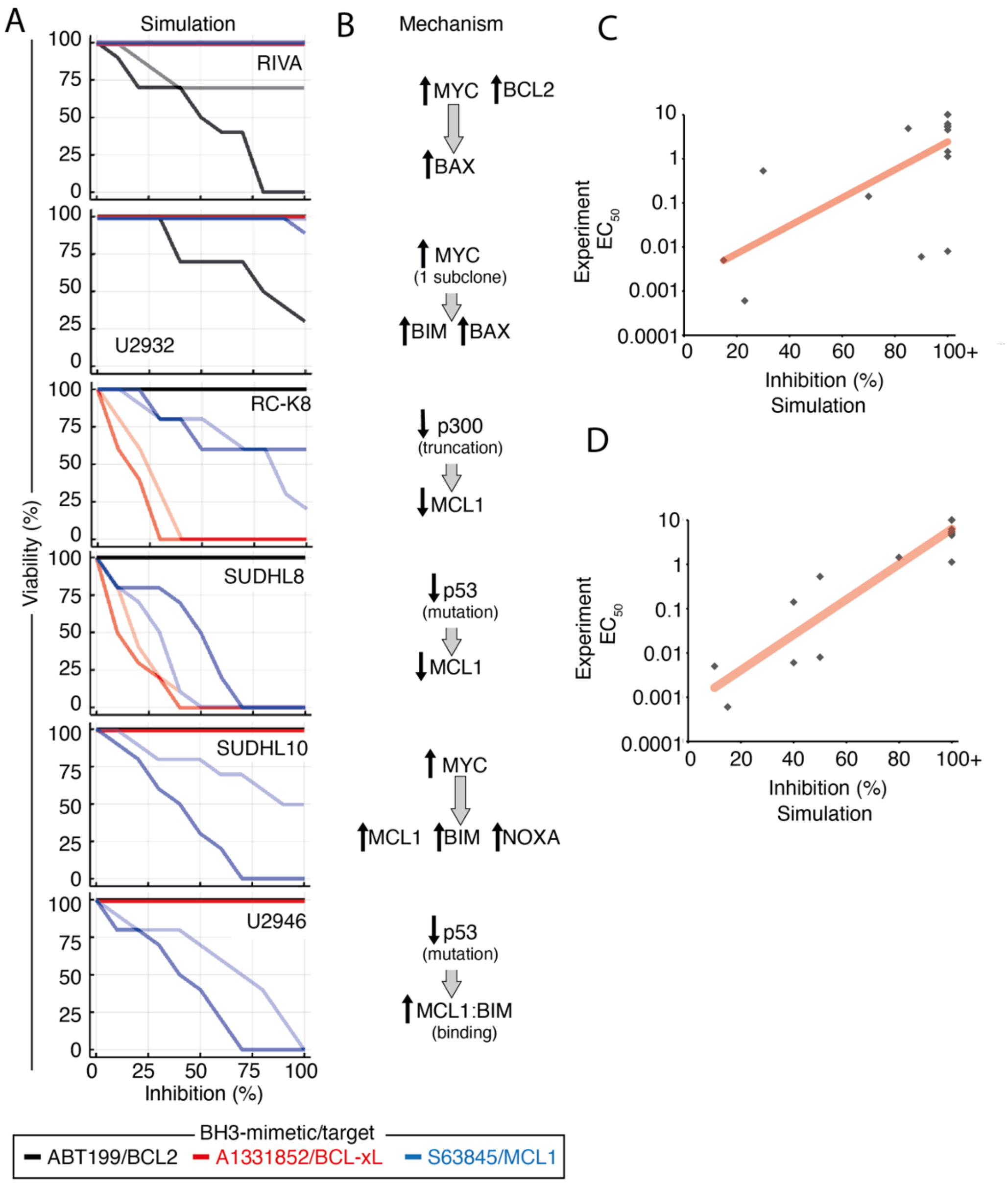


**Figure S4.** (A) The viability (%) in response to inhibition of the indicated BCL2 family proteins for the respective virtual cell lines (Figure 3) modified to consider the mutational profile of the cells. The original survival curves from Figure3 informed by protein expression alone are indicated with faint lines. Black, red and blue curves is the response corresponding to BCL2, BCL-xL and MCL1 inhibition, respectively. (B) The proposed mechanism that was simulated to improve the match between experimentally measured $EC_{50}$ values and the model prediction. (C and D) Computationally simulated inhibition percentage where 50% viability is predicted, compared to experimentally measured $EC_{50}$ values in virtual cell lines (C, $R^2$=0.38) and virtual cell lines modified to consider the mutational profile of the cells (D, $R^2$=0.67).

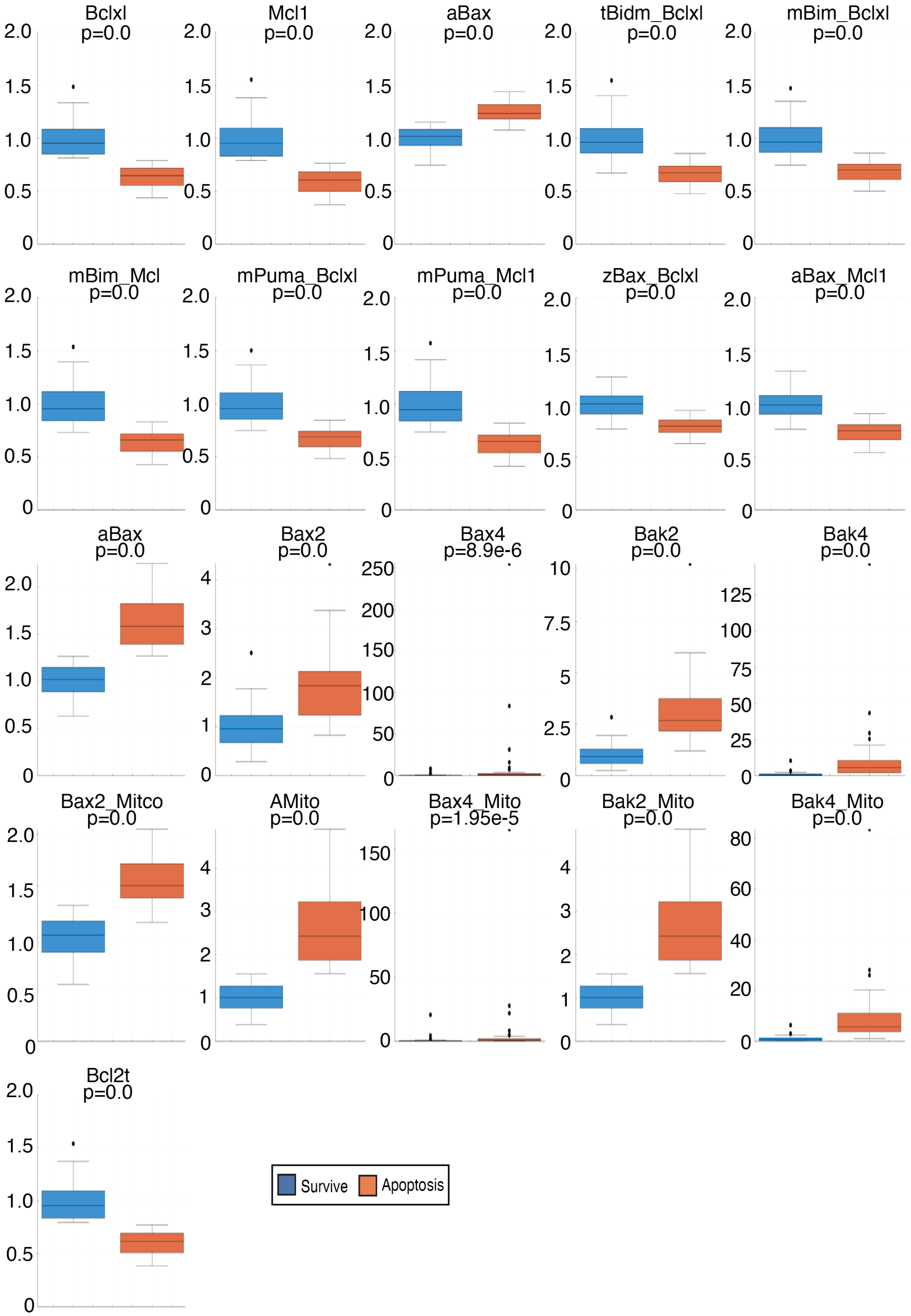


**Figure S5.** Differences in initial Protein abundances in RIVA cells between cells that survive (Blue box) and cells that undergo death (Red box).

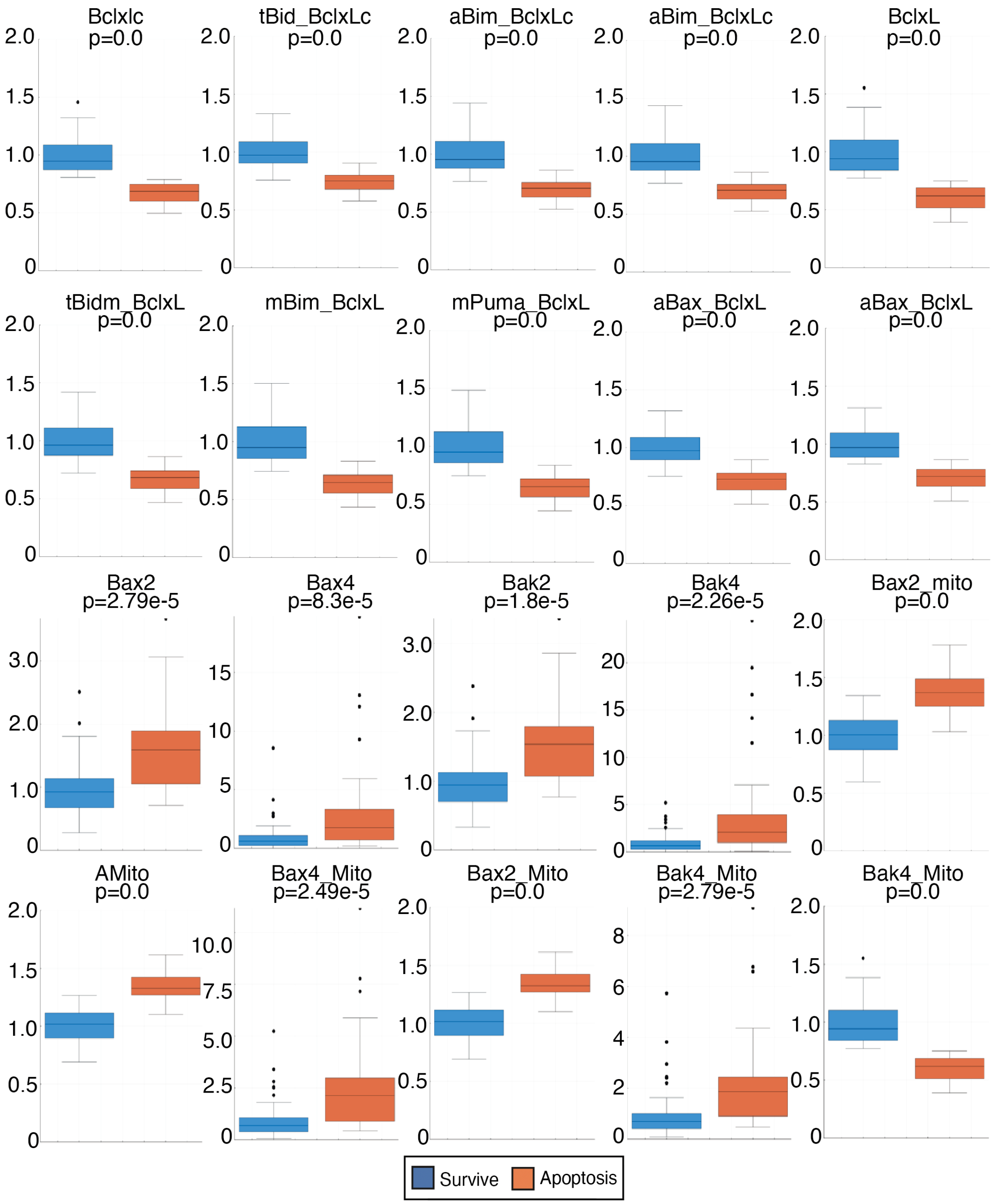


**Figure S6.** Differences in initial protein abundances in U2932 cells between cells that survive (Blue box) and cells that undergo death (Red box).

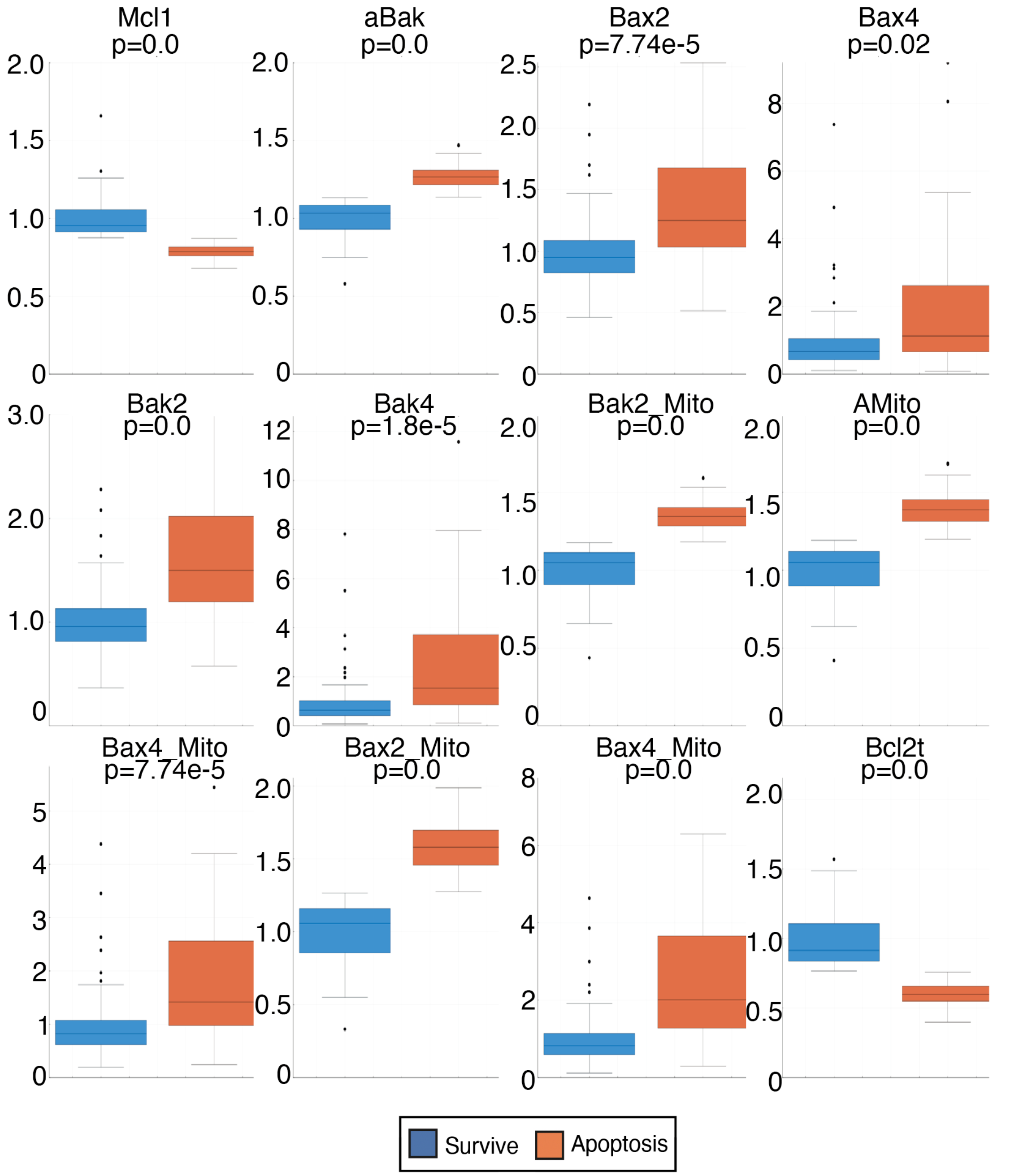


**Figure S7.** Differences in initial protein abundances in SUDHL10 cells between cells that survive (Blue box) and cells that undergo death (Red box).

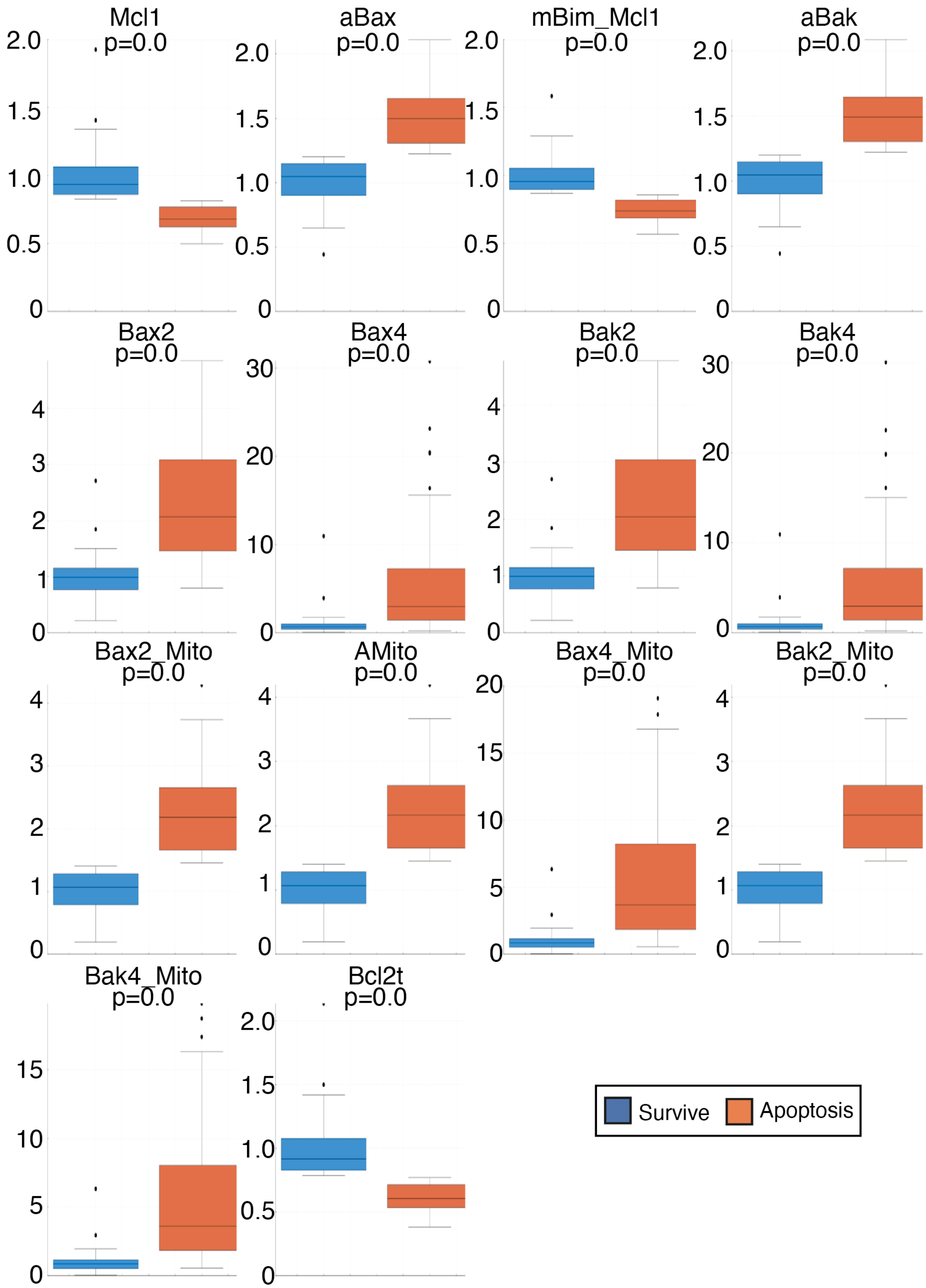


**Figure S8.** Differences in initial protein abundances in U2946 cells between cells that survive (Blue box) and cells that undergo death (Red box).

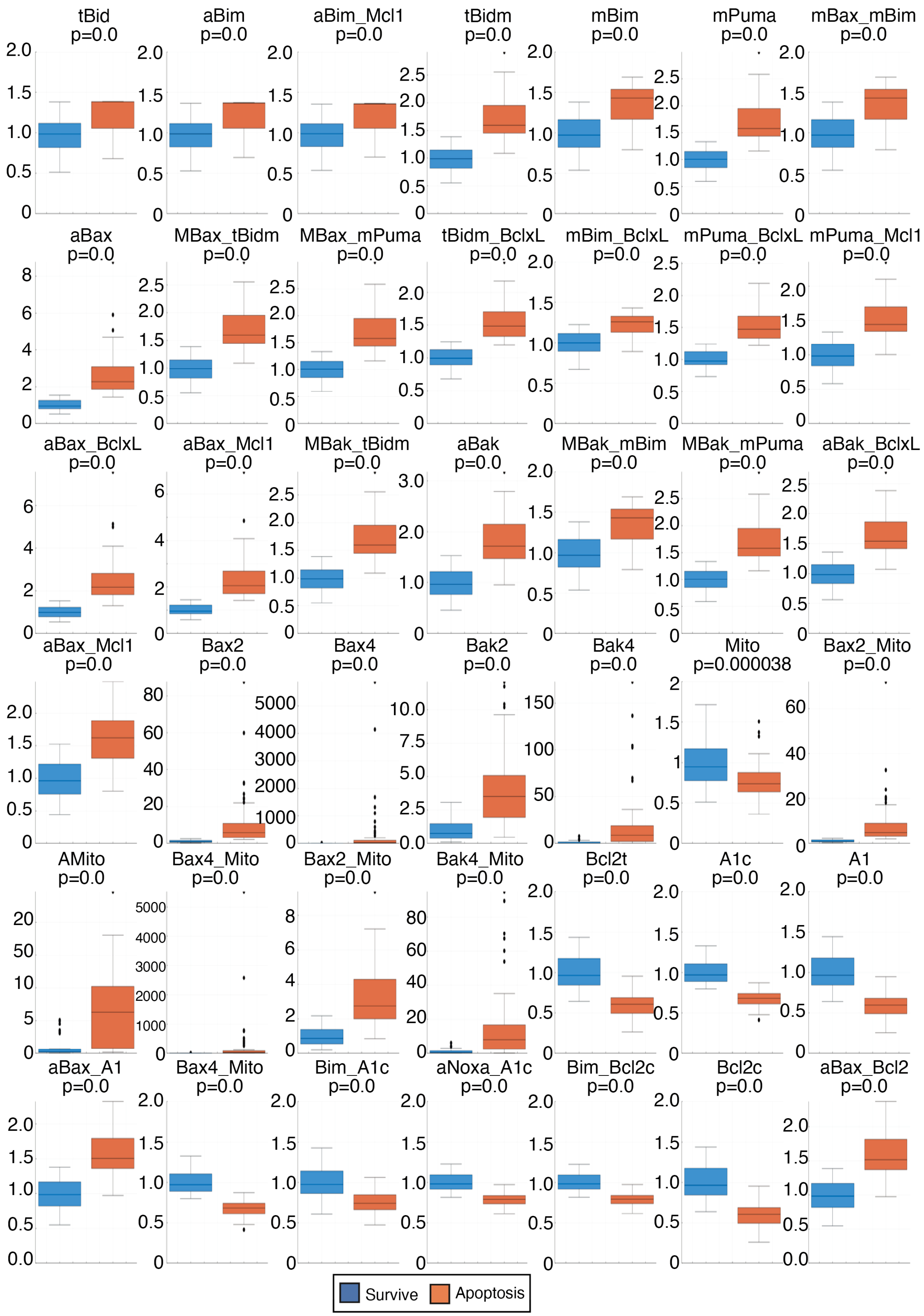


**Figure S9.** Differences in initial protein abundances in RC-K8 cells between cells that survive (Blue box) and cells that undergo death (Red box) in response to BCL-xL inhibition.

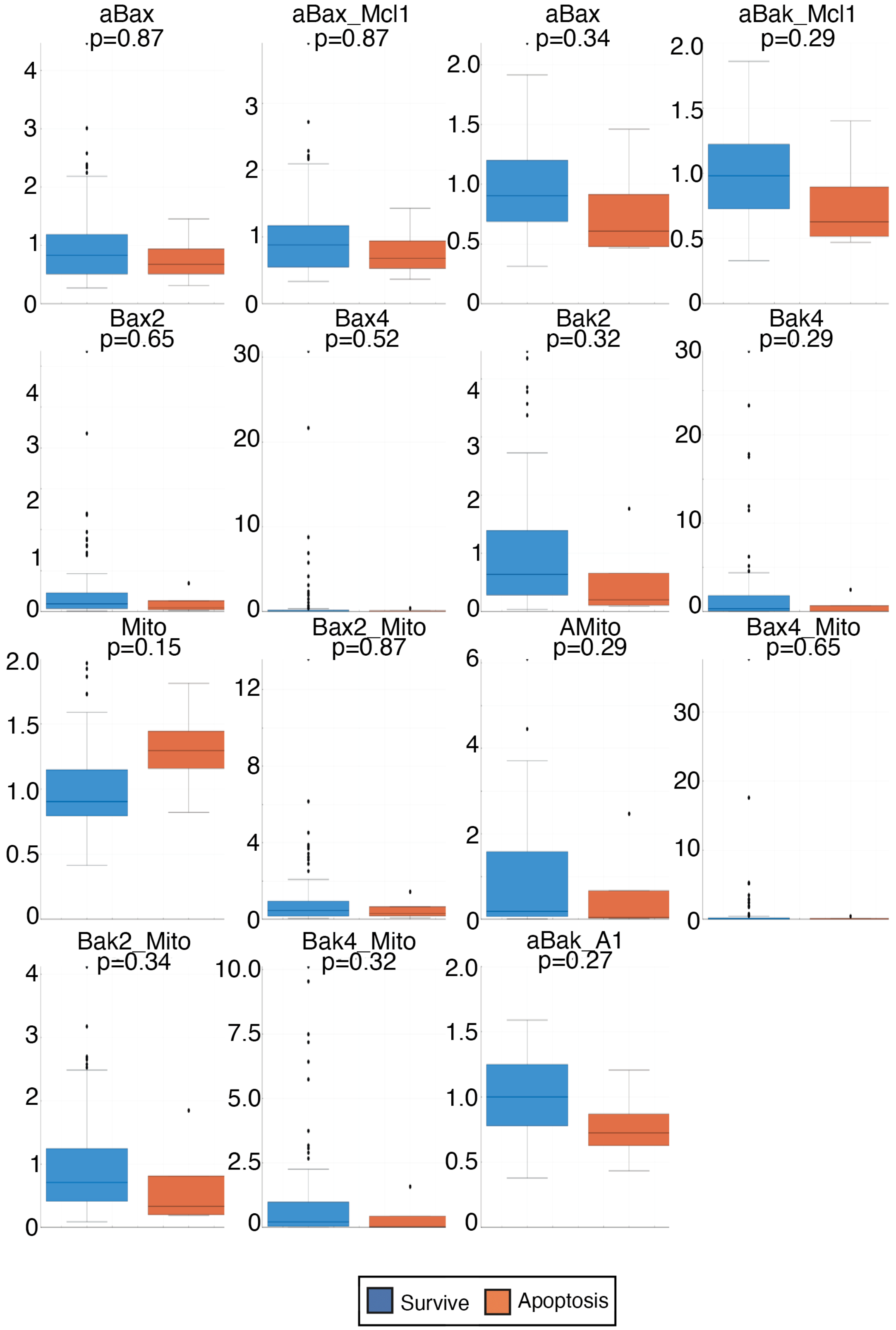


**Figure S10.** Differences in initial protein abundances in RC-K8 cells between cells that survive (Blue box) and cells that undergo death (Red box) in response to MCL1 inhibition.

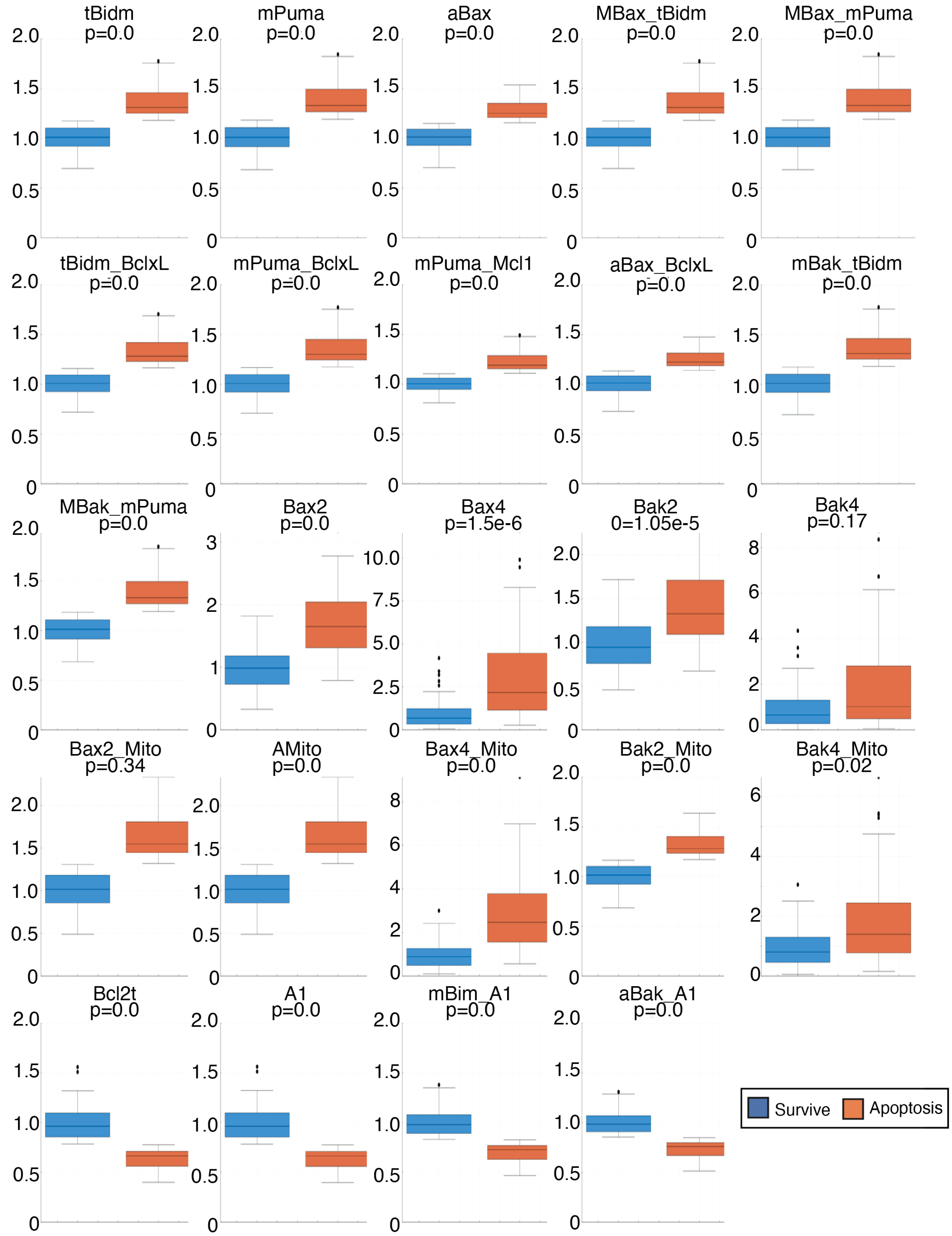


**Figure S11:** Differences in initial protein abundances in SUDHL8 cells between cells that survive (Blue box) and cells that undergo death (Red box) in response to BCL-xL inhibition.

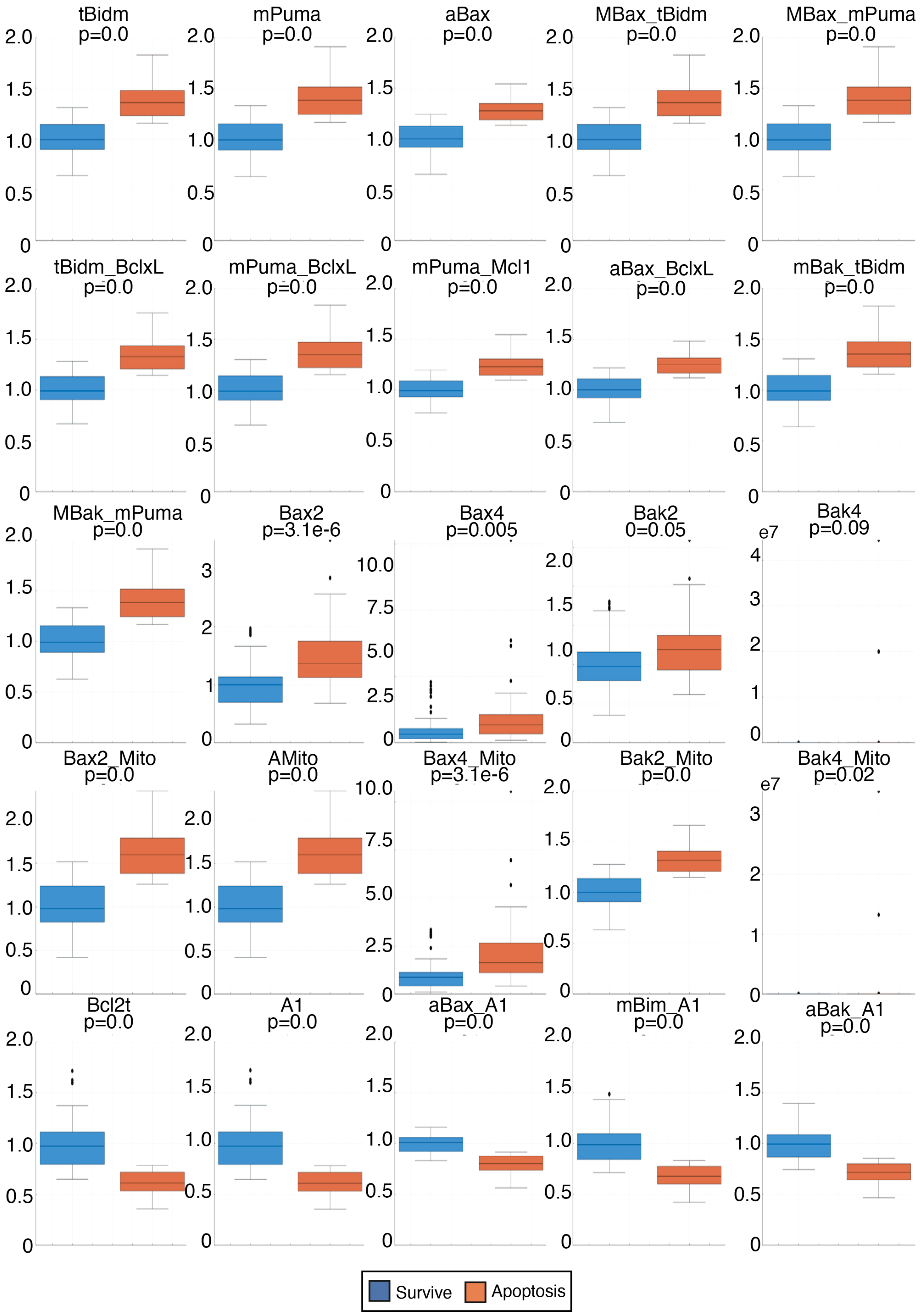


**Figure S12.** Differences in initial Protein abundances in SUDHL8 cells between cells that survive (Blue box) and cells that undergo death (Red box) in response to MCL1 inhibition.

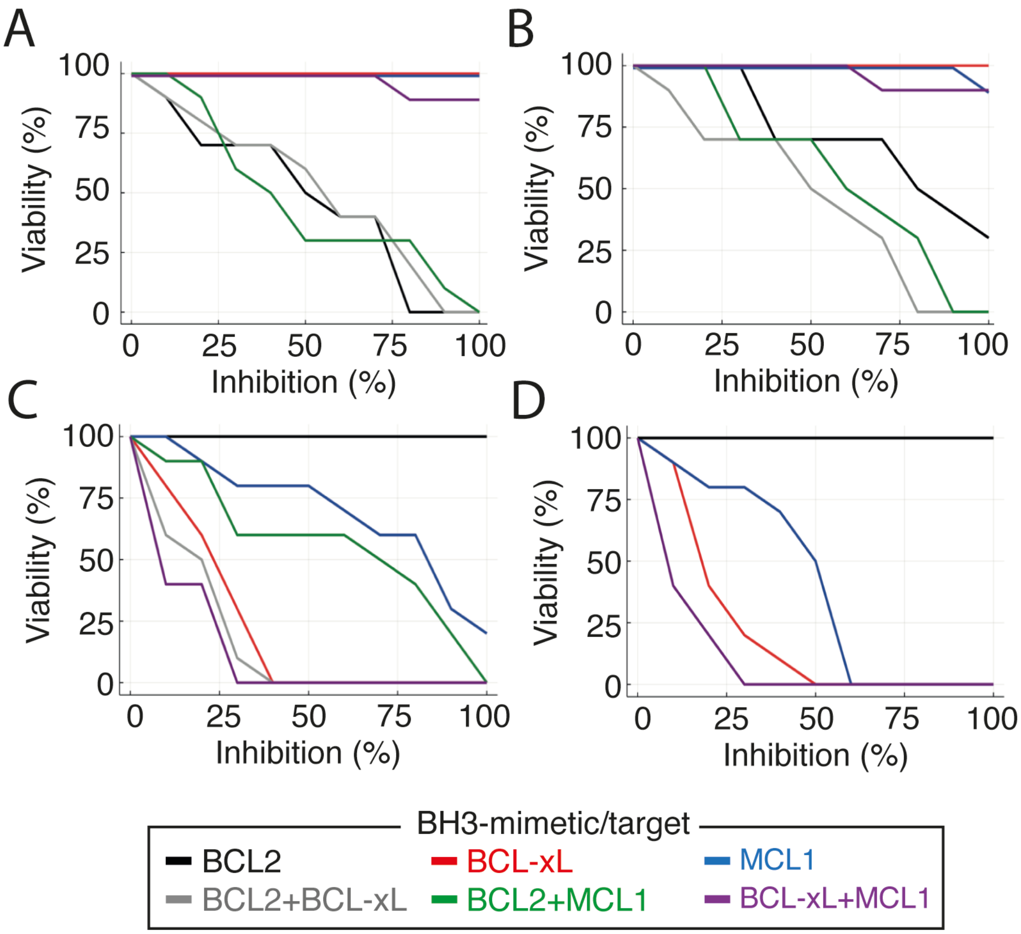


**Figure S13.** (A) Model predicts synergy between BCL2 and MCL1 inhibitors (green curve) at an intermediate range of inhibition in the RIVA cell line but no synergy between BCL2 and BCL-xL inhibitors (grey curve) and between BCL-xL and MCL1 inhibitors (purple curve). (B) In U2932, the model predicts synergy between BCL2 and BCL-xL inhibitors and between BCL2 and MCL1 but not between BCL-xL and MCL1. (C) In RC-K8, model predicts synergy between BCL-xL and MCL1 inhibitors and BCL2 and BCL-xL inhibitors but not between BCL2 and MCL1 inhibitors. (D) In SUDHL8, the model predicts synergy between BCL-xL and MCL1 inhibitors.

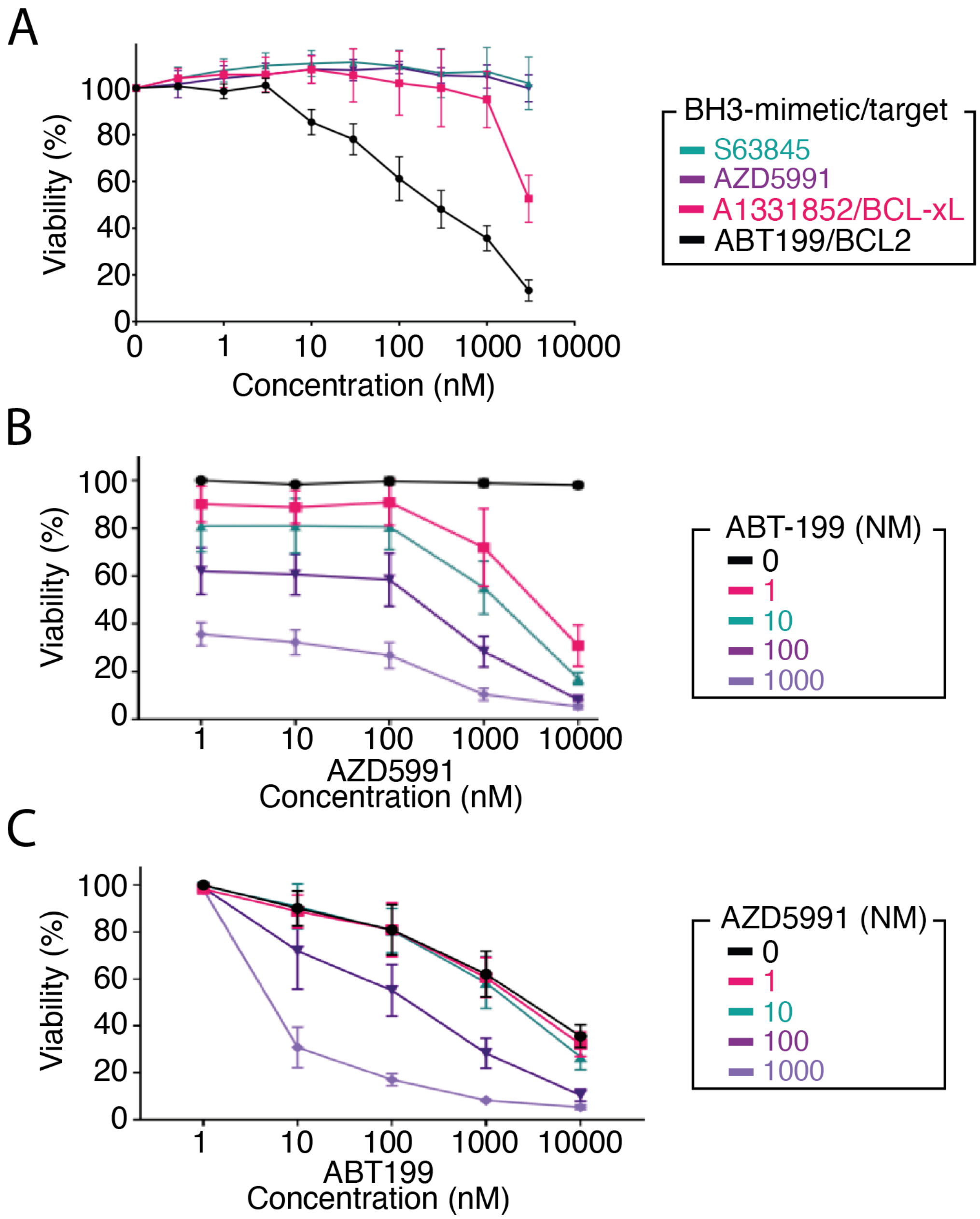


**Figure S14**. (A) Experimental data showing the % viability in response to the BH3-mimetics ABT-199 (Black curve), A1331852 (magenta curve), S63845 (teal curve) and AZD5991 (purple curve) as measured by CellTiter-Glo viability assay. (B,C) Experimental data displaying synergy induced by ABT-199 and AZD5991 combination therapy, combined into a heatmap in Figure 4.

**Table S1.** Parameters used in model

| Parameter | Value (units) | Refs. |
|---|---|---|
| BID activation | 6*e*−6 ($nM^{-1}.s^{-1}$) | [22] |
| BID inactivation | 0.06 ($s^{-1}$) | [22] |
| Active BID (tBID) | 60 (nM) | [22] |
| BIM activation | 6*e*−6 ($nM^{-1}.s^{-1}$) | [22] |
| BIM inactivation | 0.06 ($s^{-1}$) | [22] |
| Active BIM | 60 (nM) | [22] |
| PUMA activation | 1.07*e*−3 ($nM^{-1}.s^{-1}$) | [40] |
| PUMA inactivation | 1.41 ($s^{-1}$) | [40] |
| Active PUMA | 60 (nM) | based on [22] |
| NOXA activation | 1.07*e*−3 ($nM^{-1}.s^{-1}$) | [40] |
| NOXA inactivation | 1.41 ($s^{-1}$) | [40] |
| Active NOXA | 60 (nM) | based on [22] |
| tBID-BCL-xL complex association rate (cytoplasm) | 6*e*−5 ($nM^{-1}.s^{-1}$) | [22] |
| tBID-BCL-xL complex dissociation rate (cytoplasm) | 0.06 ($s^{-1}$) | [22] |
| tBID-BCL-xL complex association rate (MOM) | 6*e*−9 ($nM^{-1}.s^{-1}$) | based on [22] |
| tBID-BCL-xL complex dissociation rate (MOM) | 0.06 ($s^{-1}$) | based on [22] |
| BIM-BCL-xL complex association rate (cytoplasm) | 5.5*e*−5 ($nM^{-1}.s^{-1}$) | [28] |
| BIM-BCL-xL complex dissociation rate (cytoplasm) | 4.4*e*−4 ($s^{-1}$) | [28] |
| BIM-BCL-xL complex association rate (MOM) | 5.5*e*−7 ($nM^{-1}.s^{-1}$) | based on [28] |
| BIM-BCL-xL complex dissociation rate (MOM) | 4.4*e*−4 ($s^{-1}$) | based on [28] |
| PUMA-BCL-xL complex association rate (cytoplasm) | 8.63*e*−5 ($nM^{-1}.s^{-1}$) | [28] |
| PUMA-BCL-xL complex dissociation rate (cytoplasm) | 4.4*e*−4 ($s^{-1}$) | [28] |
| PUMA-BCL-xL complex association rate (MOM) | 8.63*e*−7 ($nM^{-1}.s^{-1}$) | based on [28] |
| PUMA-BCL-xL complex dissociation rate (MOM) | 4.4*e*−4 ($s^{-1}$) | based on [28] |
| BIM-MCL1 complex association rate (cytoplasm/MOM) | 7.8*e*−6 ($nM^{-1}.s^{-1}$) | [28] |

| | | |
|---|---|---|
| BIM-MCL1 complex dissociation rate (cytoplasm/MOM) | $2.17e^{-3}$ ($s^{-1}$) | [28] |
| PUMA-MCL1 complex association rate (cytoplasm) | $1.63e{-4}$ ($nM^{-1}.s^{-1}$) | [28] |
| PUMA-MCL1 complex dissociation rate (cytoplasm) | $5.21e^{-3}$ ($s^{-1}$) | [28] |
| PUMA-MCL1 complex association rate (MOM) | $1.63e{-6}$ ($nM^{-1}.s^{-1}$) | based on [28] |
| PUMA-MCL1 complex dissociation rate (MOM) | $5.21e^{-3}$ ($s^{-1}$) | based on [28] |
| NOXA-MCL1 complex association rate (cytoplasm) | $1.68e{-5}$ ($nM^{-1}.s^{-1}$) | [28] |
| NOXA-MCL1 complex dissociation rate (cytoplasm) | $1.39e^{-3}$ ($s^{-1}$) | [28] |
| BIM-BCL2 complex association rate (cytoplasm) | $4.48e{-5}$ ($nM^{-1}.s^{-1}$) | [28] |
| BIM-BCL2 complex dissociation rate (cytoplasm) | $1.12e^{-4}$ ($s^{-1}$) | [28] |
| BIM-BCL2 complex association rate (MOM) | $4.48e{-7}$ ($nM^{-1}.s^{-1}$) | based on [28] |
| BIM-BCL2 complex dissociation rate (MOM) | $1.12e^{-4}$ ($s^{-1}$) | based on [28] |
| PUMA-BCL2 complex association rate (cytoplasm) | $4.91e{-5}$ ($nM^{-1}.s^{-1}$) | [28] |
| PUMA-BCL2 complex dissociation rate (cytoplasm) | $1.57e^{-4}$ ($s^{-1}$) | [28] |
| PUMA-BCL2 complex association rate (MOM) | $4.91e{-7}$ ($nM^{-1}.s^{-1}$) | based on [28] |
| PUMA-BCL2 complex dissociation rate (MOM) | $1.57e^{-4}$ ($s^{-1}$) | based on [28] |
| tBID-A1 complex association rate (cytoplasm) | $8.2e{-4}$ ($nM^{-1}.s^{-1}$) | [41] |
| tBID-A1 complex dissociation rate (cytoplasm) | $1.55e^{-4}$ ($s^{-1}$) | [41] |
| tBID-A1 complex association rate (MOM) | $8.2e{-7}$ ($nM^{-1}.s^{-1}$) | [41] |
| BAX + BAX −→ BAX2 | $8.5714e^{-4}$ ($nM^{-1}.s^{-1}$) | [22] |
| BAX2 −→ BAX + BAX | 0.06 ($s^{-1}$) | [22] |
| tBID-A1 complex dissociation rate (MOM) | $1.55e^{-4}$ ($s^{-1}$) | [41] |
| BIM-A1 complex association rate (cytoplasm) | $7.8e{-4}$ ($nM^{-1}.s^{-1}$) | based on [41] |
| BIM-A1 complex dissociation rate (cytoplasm) | $3.95e^{-5}$ ($s^{-1}$) | based on [41] |
| BIM-A1 complex association rate (MOM) | $7.8e{-6}$ ($nM^{-1}.s^{-1}$) | based on [41] |
| BIM-A1 complex dissociation rate (MOM) | $3.95e^{-5}$ ($s^{-1}$) | based on [41] |

| | | |
|---|---|---|
| PUMA-A1 complex association rate (cytoplasm) | 5.9*e*−3 ($nM^{-1}.s^{-1}$) | based on [41] |
| PUMA-A1 complex dissociation rate (cytoplasm) | 5*e*−4 ($s^{-1}$) | based on [41] |
| PUMA-A1 complex association rate (MOM) | 5.9*e*−5 ($nM^{-1}.s^{-1}$) | based on [41] |
| PUMA-A1 complex dissociation rate (MOM) | 5*e*−4 ($s^{-1}$) | based on [41] |
| NOXA-A1 complex association rate (cytoplasm) | 5.9*e*−3 ($nM^{-1}.s^{-1}$) | based on [41] |
| NOXA-A1 complex dissociation rate (cytoplasm) | 1*e*−3 ($s^{-1}$) | based on [41] |
| tBID-BAX complex association rate | 6*e*−6 ($nM^{-1}.s^{-1}$) | [22] |
| tBID-BAX complex dissociation rate | 0.06 ($s^{-1}$) | [22] |
| tBID activated BAX | 60 (nM) | [22] |
| BIM-BAX complex association rate | 2.07*e*−3 ($nM^{-1}.s^{-1}$) | based on [28] |
| BIM-BAX complex dissociation rate | 0.06 ($s^{-1}$) | based on [28] |
| BIM activated BAX | 60 (nM) | based on [22] |
| PUMA-BAX complex association rate | 6*e*−6 ($nM^{-1}.s^{-1}$) | based on [28] |
| PUMA-BAX complex dissociation rate | 0.06 ($s^{-1}$) | based on [28] |
| PUMA activated BAX | 60 (nM) | based on [22] |
| tBID-BAK complex association rate | 6*e*−6 ($nM^{-1}.s^{-1}$) | [22] |
| tBID-BAK complex dissociation rate | 0.06 ($s^{-1}$) | [22] |
| tBID activated BAK | 60 (nM) | [22] |
| BIM-BAK complex association rate | 2.07*e*−3 ($nM^{-1}.s^{-1}$) | based on [28] |
| BIM-BAK complex dissociation rate | 0.06 ($s^{-1}$) | based on [28] |
| BIM activated BAK | 60 (nM) | based on [22] |
| PUMA-BAK complex association rate | 6*e*−6 ($nM^{-1}.s^{-1}$) | based on [28] |
| PUMA-BAK complex dissociation rate | 0.06 ($s^{-1}$) | based on [28] |
| PUMA activated BAK | 60 (nM) | based on [22] |
| BCL-xL-BAX complex association rate | 5.18*e*−7 ($nM^{-1}.s^{-1}$) | [28] |
| BCL-xL-BAX complex dissociation rate | 4.4*e*−4 ($s^{-1}$) | [28] |
| MCL1-BAX complex association rate | 3.25*e*−7 ($nM^{-1}.s^{-1}$) | [28] |
| MCL1-BAX complex dissociation rate | 2.6*e*−4 ($s^{-1}$) | [28] |
| BCL2-BAX complex association rate | 3.24*e*−5 ($nM^{-1}.s^{-1}$) | [28] |
| BCL2-BAX complex dissociation rate | 3.36$e^{-4}$ ($s^{-1}$) | [28] |
| A1-BAX complex association rate | 1.4*e*−6 ($nM^{-1}.s^{-1}$) | [41] |
| A1-BAX complex dissociation rate | 3.36$e^{-5}$ ($s^{-1}$) | [41] |
| BCL-xL-BAK complex association rate | 5.5*e*−6 ($nM^{-1}.s^{-1}$) | [28] |
| BCL-xL-BAK complex dissociation rate | 4.4*e*−4 ($s^{-1}$) | [28] |
| MCL1-BAK complex association rate | 3.25*e*−7 ($nM^{-1}.s^{-1}$) | [28] |
| MCL1-BAK complex dissociation rate | 2.6*e*−4 ($s^{-1}$) | [28] |
| A1-BAK complex association rate | 6.03*e*−7 ($nM^{-1}.s^{-1}$) | [41] |

| | | |
|---|---|---|
| A1-BAK complex dissociation rate | 2.6*e*−4 (s$^{-1}$) | [41] |
| BAX2 + BAX2 −→ BAX4 | 8.5714$e^{-4}$ (nM$^{-1}$.s$^{-1}$) | [22] |
| BAX4 −→ BAX2 + BAX2 | 0.06 (s$^{-1}$) | [22] |
| BAK + BAK −→ BAK2 | 8.5714$e^{-4}$ (nM$^{-1}$.s$^{-1}$) | [22] |
| BAK2 −→ BAK + BAK | 0.06 (s$^{-1}$) | [22] |
| BAK2 + BAK2 −→ BAK4 | 8.5714$e^{-4}$ (nM$^{-1}$.s$^{-1}$) | [22] |
| BAK4 −→ BAK2 + BAK2 | 0.06 (s$^{-1}$) | [22] |
| BAX2-MITO association | 8.5714 (nM$^{-1}$.s$^{-1}$) | [22] |
| BAX2-MITO dissociation | 0.06 (s$^{-1}$) | [22] |
| BAX2-induced MOMP | 60 (nM) | [22] |
| BAX4-MITO association | 8.5714 (nM$^{-1}$.s$^{-1}$) | [22] |
| BAX4-MITO dissociation | 0.06 (s$^{-1}$) | [22] |
| BAX4-induced MOMP | 60 (nM) | [22] |
| BAK2-MITO association | 8.5714 (nM$^{-1}$.s$^{-1}$) | [22] |
| BAK2-MITO dissociation | 0.06 (s$^{-1}$) | [22] |
| BAK2-induced MOMP | 60 (nM) | [22] |
| BAK4-MITO association | 8.5714 (nM$^{-1}$.s$^{-1}$) | [22] |
| BAK4-MITO dissociation | 0.06 (s$^{-1}$) | [22] |
| BAK4-induced MOMP | 60 (nM) | [22] |
| MOMP −→ MITO | 1.155245$e^{-1}$ (s$^{-1}$) | [22] |
| BAX −→ MOM | 1.16$e^{-2}$ (s$^{-1}$) | [28] |
| BAX −→ cytoplasm | 1.16$e^{-1}$ (s$^{-1}$) | [28] |
| tBID −→ MOM | 1.16$e^{-2}$ (s$^{-1}$) | based on [28] |
| tBID −→ cytoplasm | 1.16$e^{-3}$ (s$^{-1}$) | [28] |
| BIM −→ MOM | 1.16$e^{-2}$ (s$^{-1}$) | [28] |
| BIM −→ cytoplasm | 1.16$e^{-3}$ (s$^{-1}$) | [28] |
| PUMA −→ MOM | 1.16$e^{-2}$ (s$^{-1}$) | [28] |
| PUMA −→ cytoplasm | 1.16$e^{-3}$ (s$^{-1}$) | [28] |
| BCL-xL −→ MOM | 1.155245$e^{-2}$ (s$^{-1}$) | based on [22] |
| BCL-xL −→ cytoplasm | 1.155245$e^{-3}$ (s$^{-1}$) | based on [22] |
| MCL1 −→ MOM | 1.155245$e^{-2}$ (s$^{-1}$) | based on [22] |
| MCL1 −→ cytoplasm | 1.155245$e^{-3}$ (s$^{-1}$) | based on [22] |
| BCL2 −→ MOM | 1.155245$e^{-2}$ (s$^{-1}$) | [22] |
| BCL2 −→ cytoplasm | 1.155245$e^{-3}$ (s$^{-1}$) | [22] |
| A1 −→ MOM | 1.155245$e^{-2}$ (s$^{-1}$) | based on [22] |
| A1 −→ cytoplasm | 1.155245$e^{-3}$ (s$^{-1}$) | based on [22] |
| Apoptotic signal expression | 6.11505$e^{-2}$ (nM.s$^{-1}$) | [22] |
| Apoptotic signal degradation | 1.44406$e^{-3}$ (s$^{-1}$) | [22] |
| BID expression | *cell line-specific* | this work |
| BID degradation | 1.155245$e^{-2}$ (s$^{-1}$) | [22] |

| | | |
|---|---|---|
| BIM expression | *cell line-specific* | this work |
| BIM degradation | $6.91e^{1}$ ($s^{-1}$) | [28] |
| PUMA expression | *cell line-specific* | this work |
| PUMA degradation | $6.8e^{-1}$ ($s^{-1}$) | [28] |
| NOXA expression | *cell line-specific* | this work |
| NOXA degradation | $9.36e^{-1}$ ($s^{-1}$) | this work |
| BCL-xL expression | *cell line-specific* | this work |
| BCL-xL degradation | $1.155245e^{-2}$ ($s^{-1}$) | [22] |
| MCL1 expression | *cell line-specific* | this work |
| MCL1 degradation | $1.155245e^{-2}$ ($s^{-1}$) | [22] |
| BCL2 expression | *cell line-specific* | this work |
| BCL2 degradation | $6.912e^{-1}$ ($s^{-1}$) | [22] |
| A1 expression | *cell line-specific* | this work |
| A1 degradation | $9.36e^{-1}$ ($s^{-1}$) | this work |
| BAX expression | *cell line-specific* | this work |
| BAX degradation | $1.155245e^{-2}$ ($s^{-1}$) | [22] |
| BAK expression | *cell line-specific* | this work |
| BAK degradation | $6.82e^{-2}$ ($s^{-1}$) | [22] |
| tBID-BCL-xL degradation (cytoplasm/MOM) | $1.155245e^{-2}$ ($s^{-1}$) | based on [22] |
| BIM-BCL-xL degradation (cytoplasm/MOM) | $9.93e^{-1}$/$9.75e^{-1}$ ($s^{-1}$) | [28] |
| PUMA-BCL-xL degradation (cytoplasm/MOM) | $6.75e^{-2}$ ($s^{-1}$) | [28] |
| BIM-MCL1 degradation (cytoplasm/MOM) | $9.93e^{-1}$ ($s^{-1}$) | [28] |
| PUMA-MCL1 degradation (cytoplasm/MOM) | $6.93e^{-2}$ ($s^{-1}$) | [28] |
| NOXA-MCL1 degradation | $3.75e^{-2}$ ($s^{-1}$) | based on [28] |
| BIM-BCL2 degradation (cytoplasm/MOM) | $6.93e^{-1}$/$6.75e^{-1}$ ($s^{-1}$) | [28] |
| PUMA-BCL2 degradation (cytoplasm/MOM) | $6.75e^{-2}$ ($s^{-1}$) | [28] |
| tBID-A1 degradation (cytoplasm/MOM) | $6.75e^{-2}$ ($s^{-1}$) | based on [41] |
| BIM-A1 degradation (cytoplasm/MOM) | $6.93e^{-1}$/ ($s^{-1}$) | [41] |
| PUMA-A1 degradation (cytoplasm/MOM) | $6.93e^{-2}$ ($s^{-1}$) | based on [41] |
| NOXA-MCL1 degradation | $6.75e^{-2}$ ($s^{-1}$) | based on [28] |
| tBID-BAX degradation | $6.75e^{-2}$ ($s^{-1}$) | [22] |
| BIM-BAX degradation | $6.93e^{-2}$ ($s^{-1}$) | [28] |
| PUMA-BAX degradation | $6.93e^{-2}$ ($s^{-1}$) | [28] |
| tBID-BAK degradation | $6.75e^{-2}$ ($s^{-1}$) | [22] |
| BIM-BAK degradation | $6.93e^{-2}$ ($s^{-1}$) | [28] |
| PUMA-BAK degradation | $6.93e^{-2}$ ($s^{-1}$) | [28] |
| BAX-BCL-xL degradation | $1.155245e^{-2}$ ($s^{-1}$) | [28] |
| BAX-MCL1 degradation | $6.93e^{-2}$ ($s^{-1}$) | [28] |
| BAX-BCL2 degradation | $6.93e^{-2}$ ($s^{-1}$) | [22] |

| | | |
|---|---|---|
| BAX-A1 degradation | $1.155245e^{-2}$ ($s^{-1}$) | [41] |
| BAK-BCL-xL degradation | $6.93e^{-2}$ ($s^{-1}$) | [28] |
| BAK-MCL1 degradation | $6.75e^{-2}$ ($s^{-1}$) | [28] |
| BAK-A1 degradation | $6.75e^{-2}$ ($s^{-1}$) | [41] |
| BID degradation (cytoplasm/MOM) | $1.155245e^{-2}$ ($s^{-1}$) | [22] |
| BIM degradation (cytoplasm/MOM) | $6.91e^{-1}$ ($s^{-1}$) | [28] |
| PUMA degradation (cytoplasm/MOM) | $6.8e^{-1}$ ($s^{-1}$) | [28] |
